# Targeting PKC alleviates iron overload in diabetes and hemochromatosis

**DOI:** 10.1101/2023.11.28.569107

**Authors:** Somesh Banerjee, Shaolei Lu, Anand Jain, Irene Wang, Hui Tao, Shanthi Srinivasan, Elizabeta Nemeth, Peijian He

## Abstract

Diabetes is one of the most prevalent chronic diseases worldwide. Iron overload increases the incidence of diabetes and aggravates diabetic complications that cause mortality. Reciprocally, diabetes potentially promotes body iron loading, but the mechanism remains not well understood. In this study, we demonstrated systemic iron excess and the upregulation of iron exporter ferroportin (Fpn) in the enterocytes and macrophages of multiple diabetic mouse models. Increased Fpn expression and iron efflux was also seen in the enterocytes of type 2 diabetic human patients. We further showed that protein kinase C (PKC), which is activated in hyperglycemia, was responsible for the sustained membrane expression of Fpn in physiological and in diabetic settings. For the first time, we identified that PKCs were novel binding proteins and positive regulators of Fpn. Mechanistically, hyperactive PKC promoted exocytotic membrane insertion while inhibited the endocytic trafficking of Fpn in the resting state. PKC also protected Fpn from internalization and degradation by its ligand hepcidin dependent on decreased ubiquitination and increased phosphorylation of Fpn. Importantly, the loss-of-function and pharmacological inhibition of PKC alleviated systemic iron overload in diabetes and hemochromatosis. Our study thus highlights PKC as a novel target in the control of systemic iron homeostasis.

## Introduction

Iron is essential for the synthesis of hemoglobin in mammals and is a critical micronutrient for cells to execute fundamental metabolic processes. Disruption of iron homeostasis can lead to iron deficiency or iron overload. In recent years, iron overload has been increasingly recognized as an important health problem. Iron in excess mediates the pathogenesis of various pathological conditions including hepatic, metabolic, cardiovascular, and neurodegenerative diseases (1, 2). Ferroportin (Fpn, also known as SLC40A1), the only iron exporter, plays a crucial role in systemic iron metabolism by mediating dietary iron absorption in the enterocytes and iron recycling by the macrophages (3). The expression and activity of Fpn is subject to negative regulation by its ligand hepcidin (4), a peptide hormone produced by the hepatocytes in response to iron overload and inflammation (5). Hepcidin acts to suppress intestinal iron absorption and macrophage-mediated iron release by occluding Fpn and inducing its ubiquitination, internalization, and degradation (4). Thus, the hepcidin-Fpn axis plays a critical role in maintaining systemic iron homeostasis. Disruption of the hepcidin-Fpn axis is commonly seen in hereditary hemochromatosis (HH), an inherited disorder featured for systemic iron overload because of mutation(s) in genes that regulate hepcidin expression or mutation(s) in Fpn that causes resistance to hepcidin (6, 7). Beyond the role of hepcidin, other mechanisms of Fpn regulation that modulates systemic iron status remain poorly understood.

Diabetes is a metabolic disorder resulting from insulin deficiency or insulin resistance, which underlie the pathogenesis of type 1 and type 2 diabetes, respectively (8). Iron accumulation in pancreatic β cells can cause the death of β cells and a deficiency in insulin production mediating the development of type 1 diabetes (9). Elevation in body iron content also increases the incidence of type 2 diabetes (1, 10, 11), and worsens diabetic complications (12). Reducing body iron levels by restricting iron intake, by phlebotomy, or by iron chelation improves insulin sensitivity (13), decreases hypertriglyceridemia (14), and restores renal function (15) in human patients. It has further been shown that a high-iron diet increases blood glucose level in type 2 diabetic db/db (*Lepr^db^*) mice (16), while a low-iron diet ameliorates diabetic nephropathy in db/db mice (17) and improves insulin sensitivity in ob/ob *(Lep^ob^)* mice (18). Despite the detrimental effects of iron overload on diabetes, diabetes feedforward promotes body iron loading. Zheng *et al*. reported an increase in hepatic iron deposition in type 2 diabetic patients as compared to the healthy controls (19). Increased iron content was also seen in the serum, liver, heart, kidney, pancreas, and vasculature in experimental models (20–23). However, the mechanisms by which the diabetic context increases body iron content are not well understood. Wang *et al*. showed that a downregulated expression of hepcidin accounts for diabetic iron overload in type 1 and 2 diabetic rats (22). Other studies, however, failed to identify the deficiency of hepcidin in diabetic rodents (24, 25). A better understanding of the iron status in diabetes and the underlying mechanism of regulation would provide new insights for developing alternative treatments for diabetic control.

The protein kinase C (PKC) family, which consists of conventional (α, β and γ), novel (δ, χ, 1, and θ) and atypical (τ and σ) isoforms, are key signaling molecules downstream of hyperglycemia and mediate a plethora of diabetes-associated pathological conditions, including neuropathy, nephropathy, retinopathy, and cardiovascular diseases (26). PKC regulates a variety of cellular functions including the desensitization of membrane receptors, structural change and trafficking of ion transporters/channels, immune responses, cell growth, learning and memory (26). The PKCs act primarily through a phosphorylation-dependent mechanism that mediates gene transcription and post-translational protein modifications (27). Our previous studies have shown that diabetes also induces PKC activation in intestinal epithelial cells (28, 29). We further showed that PKCα promotes the expression of divalent metal transporter 1 (DMT1), a non-heme iron importer, in the luminal membrane of enterocytes (29). In the current study, by utilizing multiple mouse models of diabetes and human specimens, we demonstrated for the first time that PKC plays a critical role in the upregulation of Fpn, responsible for the development of systemic iron overload in diabetes. Mechanistically, PKC regulates the endocytic and exocytotic trafficking of Fpn in resting state and in response to hepcidin. We further showed that inhibition of PKC with a pharmacological inhibitor is effective in combating against HH-associated iron overload through attenuation of Fpn expression. Collectively, our findings highlighted a novel role of PKC in Fpn regulation and systemic iron homeostasis, and that PKC signaling may be targeted to alleviate systemic iron overload related to HH or other pathological conditions.

## Results

### Diabetes increases body iron loading and Fpn expression independent of hepcidin

To determine body iron status associated with diabetes, we utilized multiple mouse models of diabetes including streptozotocin (STZ)-induced type 1 diabetic mice, and type 2 diabetic db/db and KKAy mice. STZ is a cytotoxic compound that selectively accumulates in pancreatic β-cells and causes their destruction. In the type 2 diabetic models, hyperphagia, obesity and hyperglycemia result from a mutation in the leptin receptor (db/db) or a polygenic cause (KKAy).

STZ treated mice were studied at 1 and 8 weeks after the last STZ injection that represent acute and chronic phases of diabetes. The 1-week and 8-week groups of STZ treated mice developed a comparable degree of hyperglycemia (**Fig. 1A**). Increased serum iron content was seen in both groups of STZ-treated mice as compared to the non-diabetic controls (**Fig. 1B**). Liver is the primary site of iron deposition when body iron is in excess. Indeed, a significant increase in liver non-heme iron was observed in the 8-week STZ group (**Fig. 1C**). The increase in liver iron content was recapitulated by *Perl*’s iron staining, with a stronger staining in periportal hepatocytes (**Fig. 1D**). We used db/db mice at 14 weeks of age which represents approximately 8 weeks on hyperglycemia (**Fig. 1E**). Compared to the control db/+ mice, db/db mice also displayed significant increases in serum (**Fig. 1F**) and liver iron content (**Fig. 1G**). Hepatic iron overload was confirmed by *Perl*’s iron staining. Unlike STZ-treated mice, wherein iron was primarily deposited in the hepatocytes, db/db mice exhibited iron loading in both the hepatocytes and Kupffer cells (**Fig. 1H**). Type 2 diabetic KKAy mice started to develop hyperglycemia at approximately 12 weeks and were studied at 20 weeks when blood glucose was prominently increased (**Fig. 1I**). Like the STZ-treated and db/db mice, KKAy mice also showed an elevation in serum iron compared to their controls (**Fig. 1J**). However, despite higher serum iron, liver iron content was lower in KKAy than the control mice by biochemical assay (**Fig. 1K**) and iron staining (**Fig. 1L**). We wondered whether this was due to the lesser liver iron uptake from the circulation. Transferrin receptor 1 (TfR1) and ZIP14 are key iron importers that mediate the uptake of transferrin-bound and non-transferrin-bound iron, respectively (30, 31). We found that ZIP14 expression was not different between the KKAy and control mice. However, TfR1 mRNA level was significantly lower by 45% in KKAy mice (**Fig. 1M**), which may account for the lower iron content in the liver of KKAy mice.

**Figure 1.**
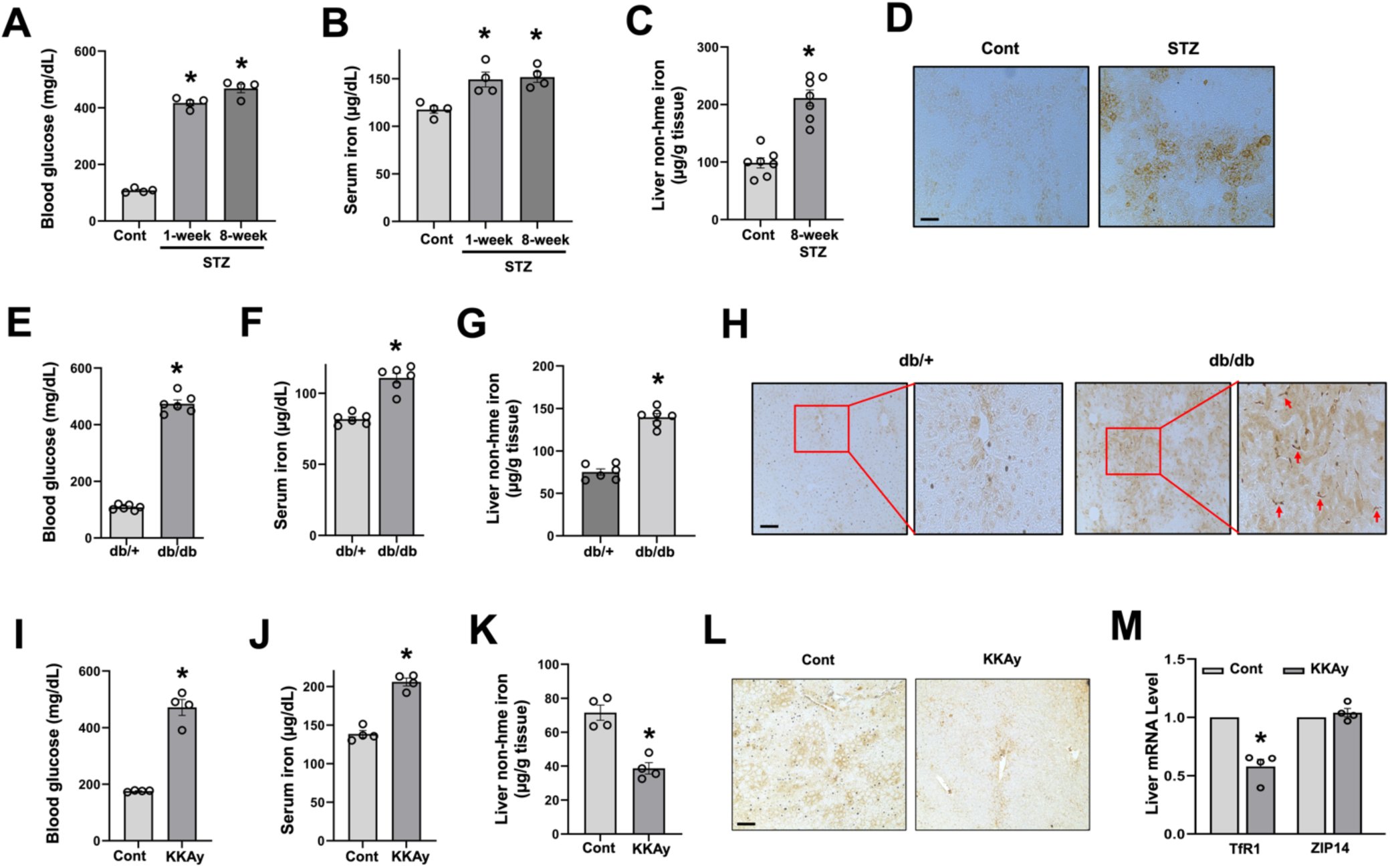
Body iron status in different diabetic mouse models. Male mice treated by streptozotocin (STZ) (**A**-**D**), male db/db mice (**E**-**H**) and KKAy mice (**I**-**M**) were studied in comparison to their corresponding controls (Cont). Blood glucose (**A**, **E**, **I**), serum iron (**B**, **F**, **J**) and liver non-heme iron (C, G, K) levels were determined. *Perl*’s iron staining was performed to show the magnitude of iron deposition in the liver of diabetic and control mice (**D**, **H**, **L**). (**M**) Hepatic TfR1 and ZIP14 mRNA expression was determined in KKAy and control mice by qRT-PCR using the *Rpl13a gene* as the internal control. Data are expressed as mean ± SEM (n = 4-7 mice *per* group). * *P* < 0.01 compared to the corresponding controls. *Red arrows* denote Kupffer cells in the liver of db/db mice. *Bars*, 100 μm.

A major mechanism that leads to an elevation in serum iron is through the upregulation of Fpn-mediated intestinal iron absorption and/or iron release from the macrophages such as in HH. We assessed Fpn expression in the duodenum, and in spleen which has abundant iron-recycling macrophages. The protein expression of Fpn was significantly increased in the duodenum (**Fig. 2A**) and spleen (**Fig. 2B**) in STZ-treated mice compared to the controls. The increase was more robust in the duodenum than the spleen (**Fig. 2A**). Of note, an increase in Fpn expression was noticed in the duodenum as early as 1 week of hyperglycemia. Same Fpn increases were seen in db/db mice as compared to the db/+ controls (**Fig. 2C** and **2D**). Upregulation of Fpn was also observed in KKAy mice, but a stronger increase was seen in the spleen (**Fig. 2E** and **2F**). We also determined the expression of F4/80, a macrophage marker, to assess whether increased Fpn in the spleen of diabetes is potentially due to a higher number of macrophages. Western blotting showed that F4/80 expression was not different between the diabetic and control mice in any of the studied models (**Fig. 2B, 2D** and **2F**).

**Figure 2.**
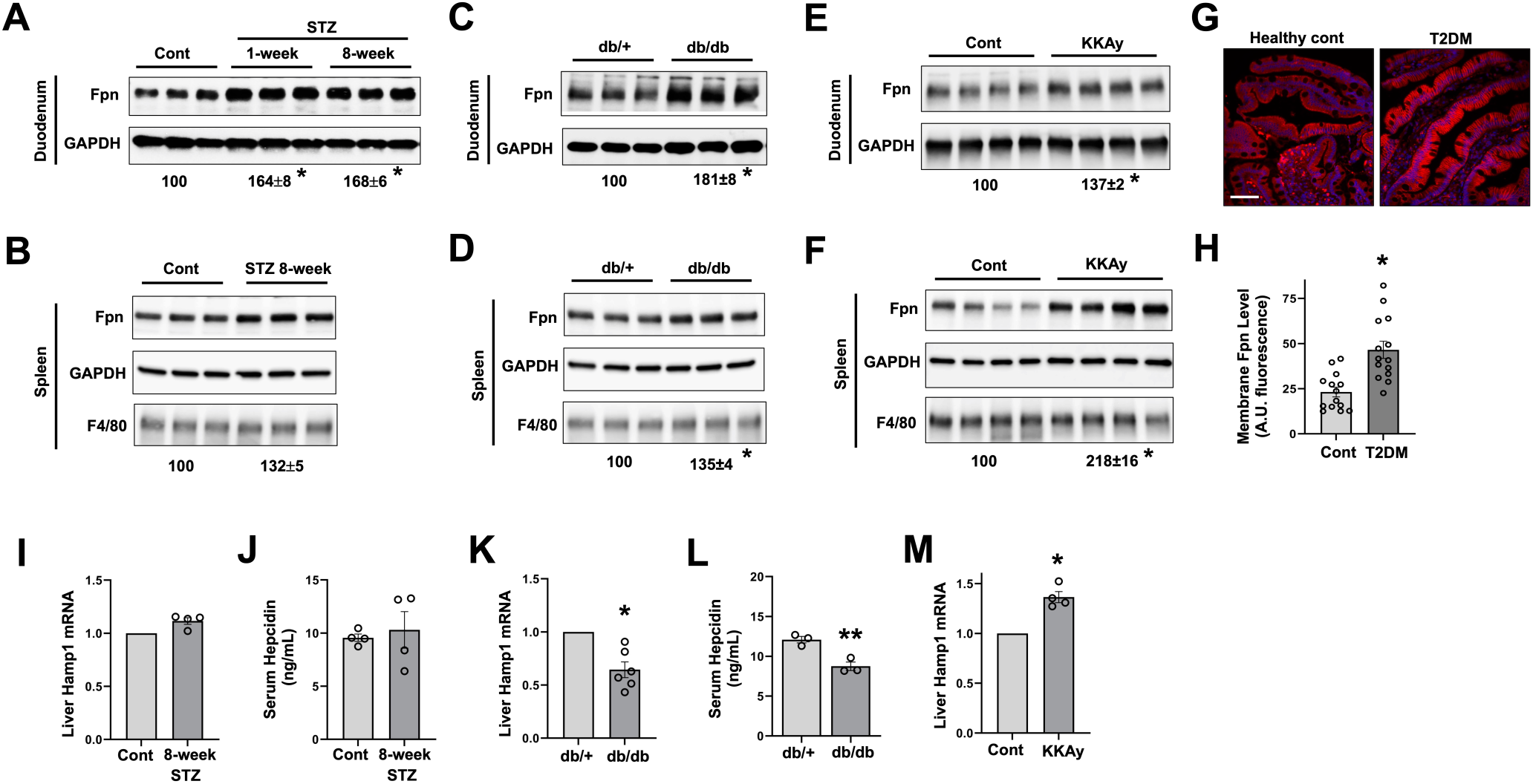
Upregulation of Fpn expression in diabetic mice and humans. Fpn protein expression in the duodenum (**A**, **C**, **E**) and spleen (**B**, **D**, **F**) of STZ-treated mice (**A**-**B**), db/db mice (**C**-**D**) and KKAy mice (**E**-**F**), with GAPDH as an internal control. The percentages of change in Fpn expression (underneath the blots) in diabetic mice over the control are shown as mean ± SEM (n = 6 for **A**-**D**, and n = 4 for **E** and **F**). (**G**) Representative Fpn staining of the duodenum of type 2 diabetic (T2DM) and non-diabetic human subjects. (**H**) Quantified arbitrary fluorescent intensity of membrane Fpn in T2DM and control subjects. Data are mean ± SEM (n = 14 human subjects). For each duodenal sample, fluorescent signal was measured in 10 representative cells by ImageJ and the average was obtained. Liver Hamp1 mRNA expression was determined by qRT-PCR in STZ-treated mice (**I**), db/db (**K**) and KKAy (**M**) mice. Serum hepcidin content was determined in STZ-treated (**J**) and db/db (**L**) mice by ELISA. * *P* < 0.01, and ** *P* < 0.05 compared to the corresponding controls. *Bar*, 50 μm.

Next, we determined whether Fpn expression is altered in diabetic human patients, with a focus on type 2 diabetes given its high prevalence. Duodenal specimens of diabetic and non-diabetic human subjects, both men and women, were collected from a pathology archive. Subjects were excluded if they have duodenal inflammation, hereditary hemochromatosis, and are on insulin therapy or iron supplementation. Women of perimenopausal age (45-55 years old) were also excluded due to potentially increased expression of hepcidin associated with menopause (32, 33). We also excluded patients with severe anemia but included six diabetic subjects with mild to moderate anemia (hemoglobin level in the range of 10-12 g/dL), because it is difficult to avoid involving anemic subjects given the high prevalence of anemia in diabetic patients (34, 35). The expression and localization of Fpn in the enterocytes varied among the individuals in both the diabetic and control groups (**Fig. 2G**, and **Fig. S1**). By quantifying the fluorescence intensity of Fpn staining, we concluded that membrane Fpn expression was significantly increased in diabetic subjects relative to the controls (**Fig. 2H**). Moreover, iron sequestration was seen in the enterocytes of control subjects but was rarely observed in diabetic patients (**Fig. S2**). These findings in mice and humans demonstrate that intestinal Fpn is upregulated accounting for diabetes-associated body iron overload.

Hepcidin is a master negative regulator of Fpn (4). A previous study showed that liver hepcidin expression was decreased in STZ-induced type 1 diabetic rats (22). We failed to see a significant decrease in hepatic hepcidin mRNA (**Fig. 2I**) or serum hepcidin content (**Fig. 2J**) in STZ-induced diabetic mice. Liver hepcidin expression (**Fig. 2K**) and serum hepcidin level (**Fig. 2L**) were significantly lower in db/db mice, which may be attributable to the impaired leptin receptor-mediated signaling (36, 37). Moreover, hepcidin mRNA expression was significantly increased by approximately 30% in KKAy mice (**Fig. 2M**). Collectively, these findings show that hepcidin expression is divergently regulated, and thus is less likely the primary mechanism responsible for the upregulation of Fpn in different models of diabetes.

### Deficiency in PKCα reduces Fpn expression alleviating diabetes-associated iron overload

Hyperactive PKC, a key signaling associated with hyperglycemia, mediates the pathogenesis of many diabetic complications (26). PKCα and PKCδ are the most abundant PKC isoforms in the intestine (28). We have previously shown that intestinal PKCα, but not PKCδ, is phosphorylated or activated in STZ-induced diabetic mice (23). We first showed that, in the physiological setting, deficiency in PKCα led to a significant decrease in serum iron content in both female and male (**Fig. 3A**). Consistently, the expression of Fpn protein was significantly decreased in the duodenum (**Fig. 3B**) and spleen (**Fig. 3C**) in PKCα^-/-^ mice. Confocal imaging further showed a decreased membrane but increased cytoplasmic pool of Fpn in PKCα^-/-^ mice (**Fig. 3D**). Defects in Fpn expression would lead to an impaired iron efflux by the enterocytes. This is confirmed by the findings of intestinal epithelial iron sequestration (**Fig. S3A**) and increased expression of ferritin heavy chain 1 (FTH1) (**Fig. S3B**), an iron binding protein and marker of iron overload, in PKCα^-/-^ mice.

**Figure 3.**
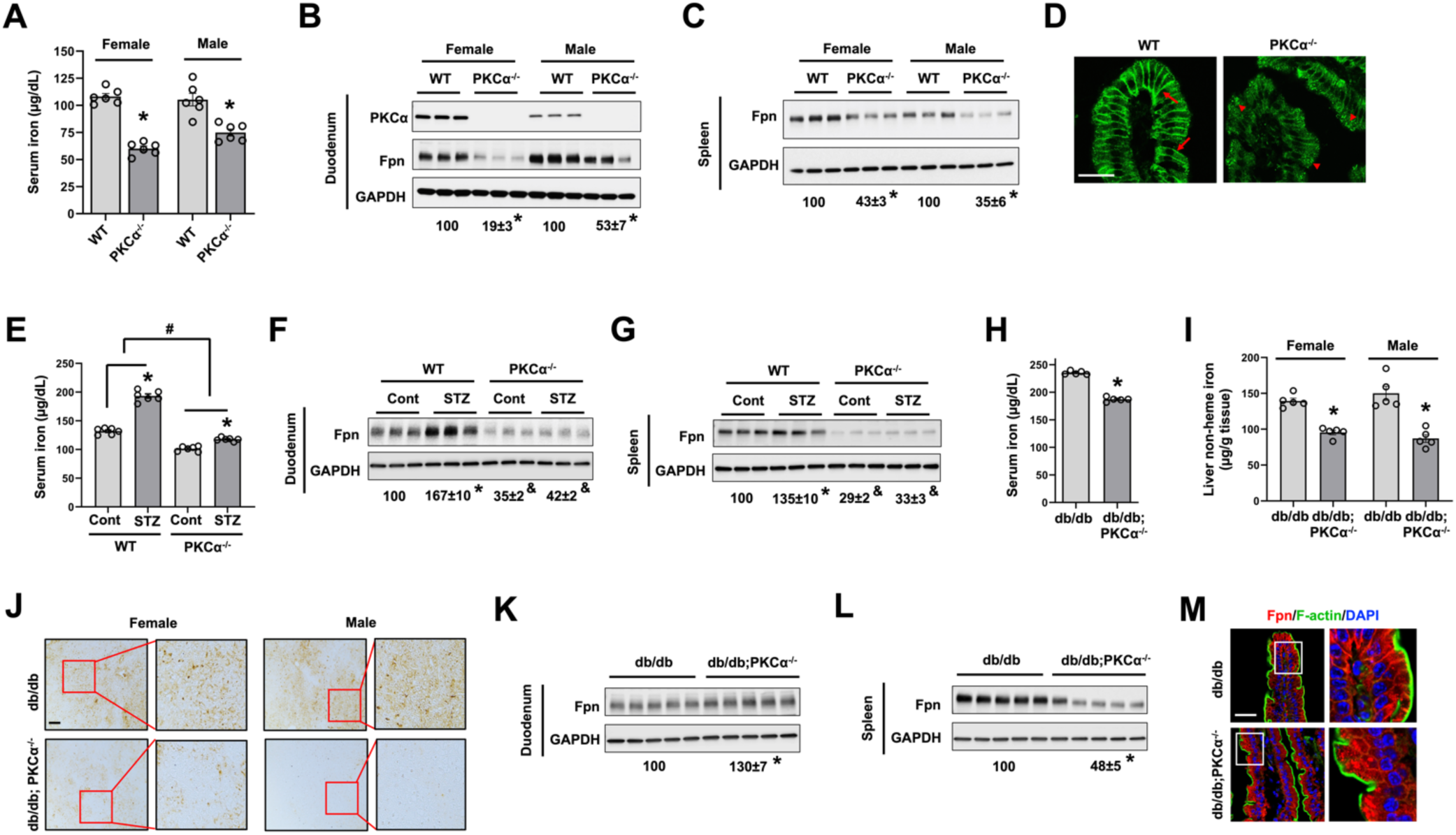
Deficiency in PKCα downregulates Fpn expression in physiology and diabetes. Serum iron (**A**), and Fpn expression in the duodenum (**B**) and spleen (**C**) were determined in 16-week-old female and male PKCα^-/-^ versus WT mice. Immunoblotting for PKCα shows the successful deletion of PKCα in PKCα^-/-^ mice. (**D**) Representative confocal images show subcellular localization of Fpn in the duodenum of PKCα^-/-^ and WT male mice. *Arrow*, membrane Fpn expression; *arrowhead*, intracellular Fpn. Serum iron (**E**), and Fpn expression in the duodenum (**F**) and spleen (**G**) were determined in WT and PKCα^-/-^ male mice with and without STZ induced diabetes. (**H**) Serum iron was determined in female db/db and db/db;PKCα^-/-^ mice. Liver iron content was determined in db/db and db/db;PKCα^-/-^ mice by biochemical assay (**I**) and *Perl*’s iron staining (**J**). Fpn protein expression was determined in the duodenum (**K**) and spleen (**L**) of db/db and db/db;PKCα^-/-^ female mice. (**M**) Confocal images showing Fpn localization in the enterocytes of db/db and db/db;PKCα^-/-^ mice. Data are shown as mean ± SEM (n = 5-6). *, *P* < 0.01 compared with their corresponding controls. #, *P* < 0.01 comparing the percentages of differences between WT and PKC^-/-^ mice. &, *P* < 0.01 compared to the WT-Cont group. *Bars*, 50 μm.

Next, we asked whether PKCα plays an important role in iron transport in diabetes. The loss-of-function of PKCα greatly attenuated diabetes-induced increase in serum iron in STZ treated mice (**Fig. 3E**), in line with our previous finding of the decreased liver iron loading in STZ-treated PKCα^-/-^ mice (29). Knockout of PKCα also blunted diabetes induced upregulation of Fpn in the duodenum (**Fig. 3F**) and spleen (**Fig. 3G**). Moreover, knockout of PKCα led to a significant reduction in serum (**Fig. 3H**) and liver iron content (**Fig. 3I** and **3J**) in type 2 diabetic db/db mice. The differences in serum and liver iron were not associated with different levels of hyperglycemia, as blood glucose level was not significantly different between the db/db and db/db;PKCα^-/-^ mice (**Fig. S4**). To our surprise, Fpn protein expression in the duodenum was significantly increased in db/db;PKCα^-/-^ mice (**Fig. 3K**), although an expected decrease was seen in the spleen (**Fig. 3L**). Through confocal microscopic analysis, we identified that Fpn in db/db;PKCα^-/-^ mice was primarily found in the cytoplasm but not on the membrane in the enterocytes (**Fig. 3M**). It is unclear whether the mislocalization of Fpn resulted from inappropriate trafficking of nascent Fpn to the cell surface, or impaired degradation of internalized Fpn in the db/db background.

Collectively, these results highlight PKCα as a previously unappreciated positive regulator of Fpn, and that hyperglycemia-mediated activation of PKCα accounts, at least in part, for diabetic iron overload through the upregulation of Fpn.

### Inhibition of PKCα promotes endocytosis and impairs exocytosis of Fpn

Next, we sought to determine the mechanism of Fpn downregulation in the loss-of-function of PKCα. Hepcidin is a master negative regulator of Fpn expression (4). Therefore, we first assessed whether the knockout of PKCα increases hepcidin. Surprisingly, hepcidin mRNA expression was not altered in male and only slightly increased in female PKCα^-/-^ mice (**Fig. S5**), suggesting that hepcidin is not the primary regulator of the downregulated Fpn expression in PKCα^-/-^ mice, which was seen in both male and female mice. To ascertain a potentially direct effect of PKCα on Fpn expression, we utilized the bone marrow-derived macrophages (BMDMs) model. Intriguingly, Fpn protein expression was significantly lower in PKCα^-/-^ BMDMs compared to the WT BMDMs (**Fig. 4A**). To exclude the potential effect that the genetic loss of PKCα imposes on BMDM differentiation, we first let the WT BMDMs undergo differentiation in M-CSF deficient medium, and then subjected the differentiated BMDMs for a treatment by Go6976, an inhibitor of PKCα, or Go6983, a pan-PKC inhibitor. In the acute phase of treatment for 4 h, neither inhibitor altered Fpn expression at the total level, but both had led to a significant decrease in membrane Fpn expression (**Fig. 4B**). After a prolonged treatment for 24 h, both inhibitors significantly reduced total Fpn expression, with a more prominent decrease in membrane Fpn expression (**Fig. 4B**).

**Figure 4.**
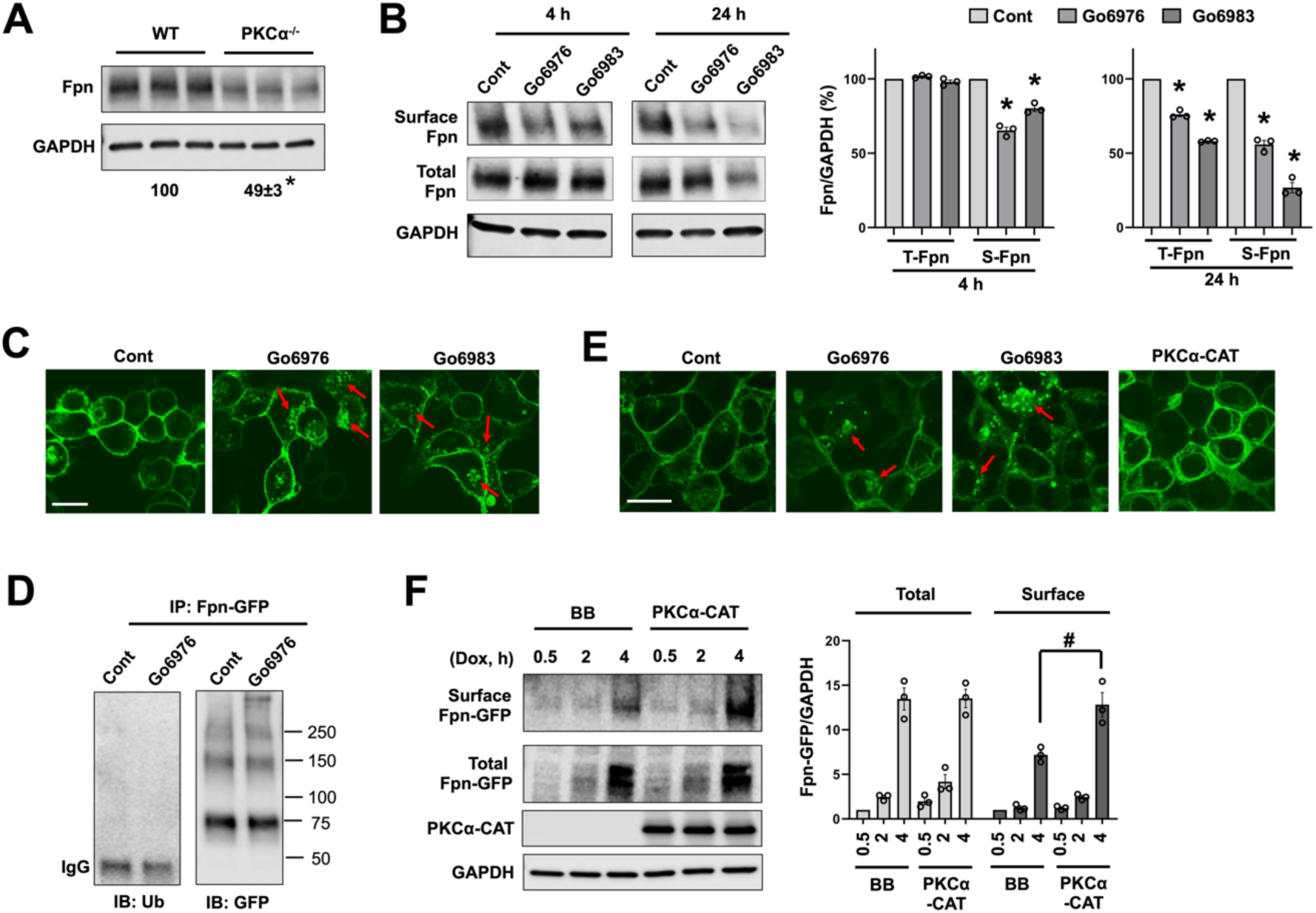
PKCα regulates the endocytic and exocytotic trafficking of Fpn. (**A**) Fpn protein expression was determined in bone marrow-derived macrophages (BMDMs) from WT and PKCα^-/-^ mice. (**B**) WT BMDMs were treated for 4 or 24 h with 5 µM Go6976 or 2 µM Go6983, followed by cell surface biotinylation assay. The total and surface Fpn expression was determined by blotting with anti-mouse Fpn antibody. (**C**) TRex-Fpn-GFP (TRex) cells were treated overnight with doxycycline (Dox, 100 ng/ml) to induce Fpn-GFP expression, followed by a 2-h pretreatment with cycloheximide (100 μg/ml), an inhibitor of *de novo* protein synthesis, prior to treating with Go6976 or Go6983 for 4 h. Subcellular localization of Fpn-GFP was then assessed by confocal microscopy. *Red arrow* denotes intracellular Fpn- GFP. (**D**) Dox induced TRex cells were treated with and without Go6976 for 4 h. Fpn was immunoprecipitated with anti-rabbit GFP antibody, and its ubiquitination level was determined by immunoblotting with anti-ubiquitin FK2 antibody. (**E**) TRex cells were transduced with PKCα-CAT, or pretreated with Go6976 or Go6983, prior to a 4-h treatment with Dox (1 µg/ml) and confocal imaging. (**F**) TRex cells transfected with backbone (BB) or PKCα-CAT were treated with Dox (1 µg/ml) for 0.5, 2 or 4 h. Surface biotinylation was performed, and the total and surface Fpn-GFP expression was determined by blotting with anti-mouse GFP antibody. Data shown are mean ± SEM of three independent experiments. * *P* < 0.01 compared to WT or the control conditions. ^#^ *P* < 0.05. *Bar*, 10 μm.

Decreased surface Fpn expression with the inhibition of PKC may result from accelerated internalization of membrane Fpn and/or reduced exocytosis of cytoplasmic Fpn. Doxycycline (Dox)-inducible HEK293:TRex-Fpn-GFP (TRex) cells were treated Dox, followed by preincubation with cycloheximide to block *de novo* protein synthesis, prior to treatments with Go6976 and Go6983. Both inhibitors induced a robust endocytosis of Fpn as revealed by the increased numbers of GFP-positive puncta in the cytoplasm (**Fig. 4C**). This process was not dependent on a change in Fpn ubiquitination (**Fig. 4D**), a known mechanism for hepcidin-induced internalization of Fpn (38). Next, we determined whether PKC modulates the exocytosis of *de novo* synthesized Fpn. TRex cells were pretreated with inhibitors of PKC or transfected with PKCα-CAT, a constitutive active form of PKCα lacking the regulatory domain (39). Cells were then treated with Dox to induce the expression of Fpn-GFP. Compared to the untreated control, cells pretreated by Go6976 and Go6983 exhibited increased expression of Fpn-GFP in the cytoplasm, with a much stronger effect seen with Go6983 (**Fig. 4E**). Conversely, cells transduced with PKCα-CAT showed a prominently higher Fpn-GFP expression in the membrane (**Fig. 4E**). The effect of PKCα on the exocytosis of Fpn was further determined by surface biotinylation. We showed that newly synthesized Fpn-GFP was delivered onto the cell surface at a much higher rate in PKCα-CAT expressing cells (**Fig. 4F**).

Collectively, these data demonstrate that PKC, including the α and possibly also other isoforms, regulates both the endocytosis and exocytosis of Fpn.

### PKCα modulates hepcidin-induced ubiquitination and internalization of Fpn

The above results showed that PKCα regulates the expression and localization of Fpn in the resting state. We wondered whether PKCα regulates the response of Fpn to its ligand hepcidin. By utilizing 293T cells that were transfected with Fpn-GFP along with PKCα-CAT or the backbone, we showed that hepcidin-induced reduction in membrane Fpn expression was prevented by the expression of PKCα-CAT (**Fig. 5A, *upper panel***). Of note, baseline membrane expression of Fpn-GFP was significantly higher in PKCα-CAT expressing cells than in the control cells, supporting the earlier results in TRex cells (**Fig. 4E** and **4F**). Two Fpn-GFP bands were noted in the total lysate lanes (**Fig. 5A, *middle panel***). They both represent Fpn protein as neither band was present in the backbone transfected cells (**Fig. S6A**). By comparing the pattern of Fpn-GFP bands in the total and surface pools of proteins, we recognized that the upper band represents the membrane form of Fpn, while the lower band represents the cytoplasmic or immature form of Fpn (**Fig. S6B**). Moreover, confocal microscopic analysis confirmed that expression of PKCα-CAT in 293T cells abolished hepcidin-induced internalization of Fpn-GFP (**Fig. 5B**).

**Figure 5.**
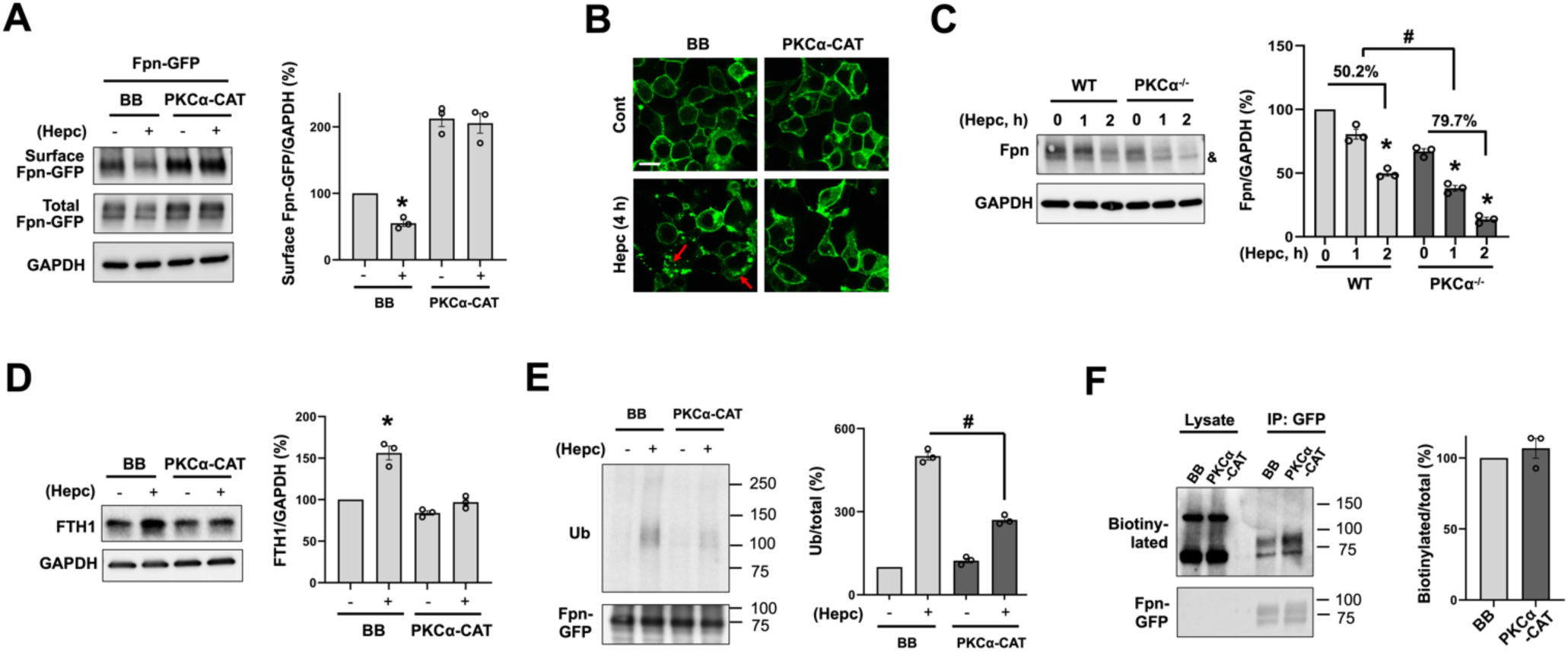
PKCα counteracts hepcidin-induced ubiquitination and internalization of Fpn. 293T cells were transfected with Fpn-GFP together with backbone (BB) or PKCα-CAT. The transfected cells were then treated for 4 h with and without 0.5 µg/ml human hepcidin (Hepc). Surface Fpn expression (**A**) and subcellular Fpn localization (**B**) were determined by surface biotinylation assay and confocal microscopy, respectively. *Red arrow* denotes intracellular Fpn-GFP. (**C**) WT and PKCα^-/-^ BMDMs were treated for 1 or 2 h with 0.25 µg/ml mouse hepcidin, and Fpn expression was determined by immunoblotting with anti-mouse Fpn antibody. & denotes a non-specific band. (**D**) 293T cells were transfected with Fpn-GFP together with BB or PKCα-CAT. The transfected cells were then treated for 24 h with and without 0.5 µg/ml human hepcidin. FTH1 expression was determined by immunoblotting. (**E**) TRex cells were transfected with BB or PKCα-CAT, and the expression of Fpn-GFP was induced overnight by 100 ng/ml Dox. Dox-induced cells were treated for 2 h with or not 0.5 µg/ml hepcidin, followed by immunoprecipitation of Fpn-GFP with anti-rabbit GFP antibody. The ubiquitination level of Fpn-GFP was determined with anti-ubiquitin FK2 antibody, and the total amount of immunoprecipitated Fpn-GFP was determined with anti-mouse GFP antibody. (**F**) Dox-treated TRex cells that were transfected with BB or PKCα-CAT were incubated for 30 min with 3 µg/ml biotinylated human hepcidin. Following immunoprecipitation with anti-rabbit GFP antibody, the abundance of co-immunoprecipitated biotinylated hepcidin was determined with HRP-conjugated streptavidin. Data are shown as mean ± SEM of three independent experiments. * *P* < 0.01 compared with the untreated controls. ^#^ *P* < 0.01. *Bar*, 10 μm.

To determine whether deficiency in PKCα sensitizes Fpn to hepcidin-induced degradation, we utilized the cultured BMDMs from WT and PKCα^-/-^ mice. Following hepcidin treatment, a significantly larger decrease in membrane Fpn expression was observed in PKCα^-/-^ BMDMs **(Fig. 5C)**. Pretreatment of 293T cells with Go6976 also potentiated hepcidin-induced internalization of Fpn (**Fig. S7**). To confirm that regulation of Fpn by PKC leads to changes in the transport activity of Fpn, we determined the expression of ferritin heavy chain 1 (FTH1), a marker of iron overload. In control cells, hepcidin induced a significant increase in FTH1 expression suggesting impaired iron efflux, whereas expression of PKCα-CAT prevented hepcidin-induced upregulation of FTH1 attributable to PKC-mediated stabilization of membrane expression of Fpn (**Fig. 5D**).

Ubiquitination of Fpn is necessary for hepcidin-induced endocytosis of Fpn (38). We observed that hepcidin-induced ubiquitination of Fpn was mitigated by the expression of PKCα-CAT (**Fig. 5E**). However, the baseline ubiquitination of Fpn was not altered (**Fig. 5E**), consistent with our earlier finding that inhibition of PKCα did not change Fpn ubiquitination level (**Fig. 4D**). One mechanism by which hyperactive PKCα attenuates the ubiquitination of Fpn could be through interfering with the binding of hepcidin to cell surface Fpn. By incubating Dox-treated cells with the biotinylated hepcidin, followed by the immunoprecipitation of Fpn-GFP, we showed that the interaction of hepcidin with Fpn was not significantly altered by the expression of PKCα-CAT (**Fig. 5F**).

Altogether, these findings demonstrate that activation of PKCα mitigates, while deficiency in PKCα potentiates, hepcidin-mediated ubiquitination and internalization of Fpn, but not through changing hepcidin-Fpn interaction.

### PKC binds and potentially phosphorylates Fpn stabilizing the membrane expression of Fpn

PKCs are protein kinase that phosphorylates substrate proteins at the serine (S) and threonine (T) residues, thereby regulating the localization and function of the substrate proteins. Confocal imaging showed that PKCα and Fpn were colocalized at the basolateral membrane in the enterocytes (**Fig. 6A**). The potential physical interaction between PKCα and Fpn was confirmed by co-IP analysis in mouse duodenum (**Fig. 6B**), and in 293T cells that express Fpn-GFP and HA-tagged full-length PKCα (**Fig. 6C**). We showed that Fpn was also bound by PKCδ (**Fig. S8A**), but whether Fpn interacts with other PKC isoforms was not examined in this study. Importantly, expression of PKCδ-CAT, the constitutively active form of PKC8, also protected Fpn from hepcidin-induced internalization in 293T cells (**Fig. S8B**). The protective role of PKCδ in Fpn regulation explains our earlier finding of stronger inhibitory effects of Go6983 (a pan-PKC inhibitor) over Go6976 (a PKCα inhibitor) on surface Fpn expression in BMDMs (**Fig. 4B**) and TRex cells (**Fig. 4E**).

**Figure 6.**
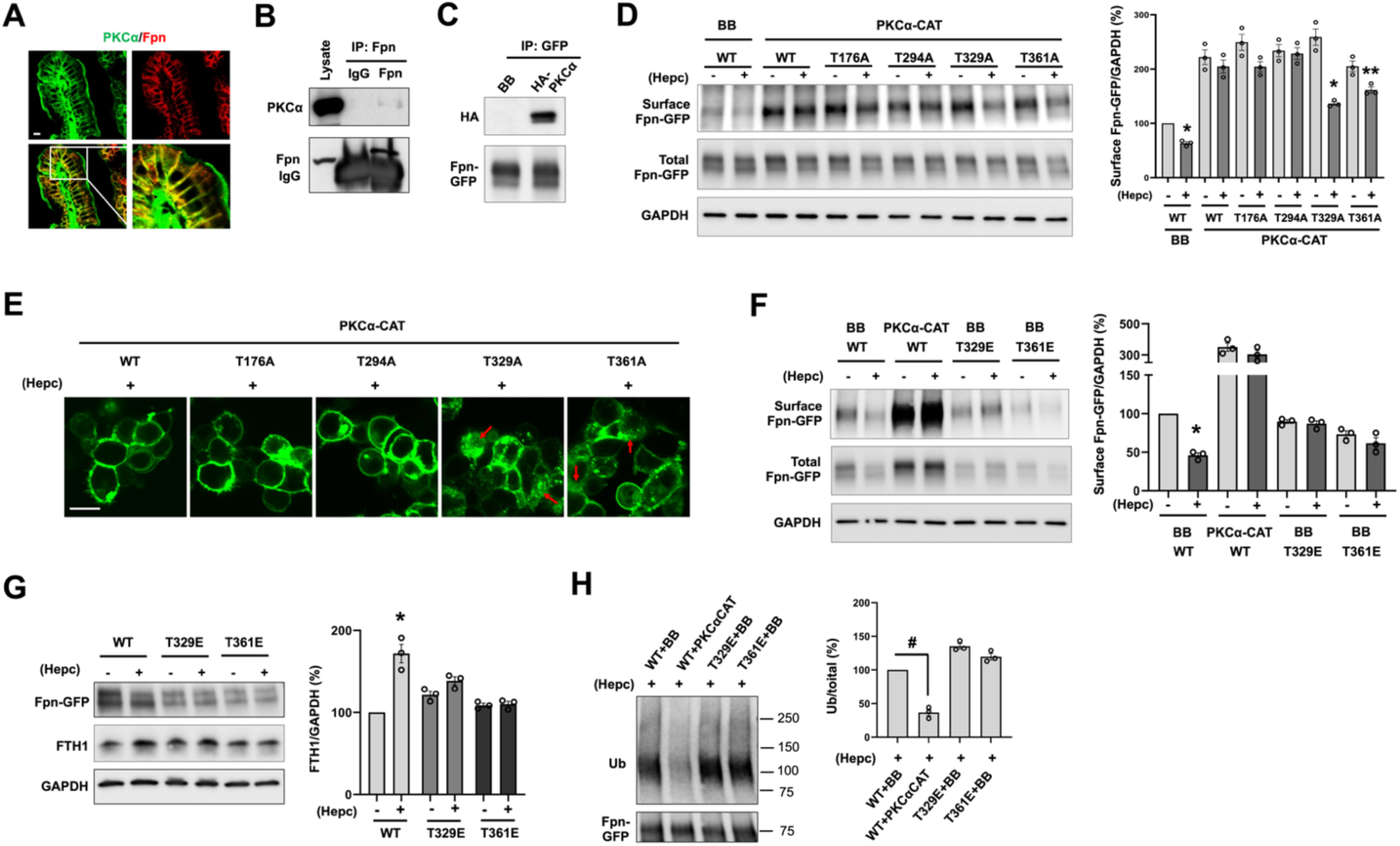
PKCα binds and potentially phosphorylates Fpn preventing hepcidin-induced internalization of Fpn. (**A**) Confocal imaging of PKCα and Fpn in the duodenum of WT mice. (**B**) Fpn was pulled down with anti-mouse Fpn antibody from mouse duodenal lysates, and the expression of PKCα was determined with rabbit anti-PKCα antibody. (**C**) Fpn-GFP was immunoprecipitated from TRex cells transfected with BB or full-length HA-PKCα, and HA-PKCα in the immunocomplex was determined with anti-HA antibody. (**D**) 293T cells were co-transfected with BB or PKCα-CAT with WT or phospho-dead mutant Fpn-GFP. Cells were treated for 4 h with and without 0.5 µg/ml hepcidin, followed by surface biotinylation assay. The cell surface and total expression of WT and mutant Fpn-GFP was determined using anti-GFP antibody. (**E**) Representative confocal images showing the localization of WT and mutant Fpn-GFP in PKCα-CAT expressing 293T cells following a 4-h treatment by hepcidin. *Red arrow* denotes intracellular Fpn-GFP. (**F**) 293T cells were co-transfected with BB or PKCα-CAT with WT, T329E or T361E mutant Fpn-GFP. Cells were subjected to surface biotinylation assay following a 4-h treatment with and without 0.5 µg/ml hepcidin. (**G**) 293T cells expressing WT, T329E or T361E mutant Fpn-GFP were incubated for 24 h with and without 0.5 µg/ml hepcidin, and the expression of Fpn-GFP and FTH1 was determined by Western blotting. (**H**) 293T cells expressing BB or PKCα-CAT with WT, T329E or T361E mutant Fpn-GFP were incubated for 2 h with or not 0.5 µg/ml hepcidin (Hepc). Fpn-GFP was immunoprecipitated with anti-rabbit GFP antibody, and its ubiquitination level was determined by immunoblotting with anti-ubiquitin FK2 antibody. The amount of the immunoprecipitated Fpn-GFP was determined with anti-mouse GFP antibody. Data are expressed as mean ± SEM of three independent experiments. * *P* < 0.01, and ** *P* < 0.05 compared to the untreated conditions. ^#^ *P* < 0.01. *Bar*, 10 μm.

By *in silico* analysis, we have identified 4 potential PKC phosphorylation sites in Fpn protein including T176, T294, T329 and T361, which are conserved in human, mouse, and many other species. The location of these threonines on the ternary structure of Fpn was predicted with AlphaFold (40) (**Fig. S9**). We then generated the phospho-dead (Thr to Ala, T to A) mutants of Fpn-GFP and assessed whether the mutants lose their protection from PKCα. Consistent with our earlier data (**Fig. 5A** and **5B**), hepcidin-mediated internalization of WT Fpn-GFP was again prevented by PKCα-CAT (**Fig. 6D**). The T176A and T294A mutants were also protected, as their membrane abundance was not altered by hepcidin treatment. In contrast, the membrane expression of the T329A and T361A mutants was significantly lower than the WT in response to hepcidin, with a more robust change in the T329A mutant (**Fig. 6D**). The loss of protection of the T329A and T361A mutants by hyperactive PKCα was further supported by confocal fluorescence imaging (**Fig. 6E**).

We then generated phosphomimetic T329E and T361E mutants to mimick hyperphosphorylated state of Fpn. Both the T329E and T361 mutant were resistant to hepcidin induced endocytosis (**Fig. 6F**). Of note, at the basal level, membrane expression of T329E (**Fig. 6F**, ***lane 5***) and T361E (**Fig. 6F**, ***lane 7***) was not higher than that of WT Fpn (**Fig. 6F**, ***lane 1***), despite that PKCα-CAT markedly increased the membrane expression of WT Fpn (**Fig. 6F**, ***lane 3***). This result implicates that phosphorylation of Fpn confers resistance of Fpn to hepcidin induced endocytosis but does not stabilize membrane Fpn expression in the resting state. The resistance of the T329E and T361E mutants was translated into sustained iron efflux in the presence of hepcidin, as indicated by the lack of FTH1 upregulation by hepcidin (**Fig. 6G**). We then asked whether T329E and T361E mutants are less ubiquitinated accounting for their resistance to hepcidin induced internalization. Surprisingly, in response to hepcidin, T329E (**Fig. 6H, *lane 3***) and T361E (**Fig. 6H, *lane 4***) showed a comparable level of ubiquitination to the WT Fpn (**Fig. 6H, *lane 1***), which was in strong contrast to the drastically decreased ubiquitination of WT Fpn in PKCα-CAT expressing cells (**Fig. 6H, *lane 2***).

These data demonstrate that PKC binds and potentially phosphorylates Fpn accounting, at least in part, for PKC-mediated protection of Fpn from the negative effect of hepcidin. However, phosphorylation of Fpn, at least on the currently studied residues, does not fully recapitulate the effects of hyperactive PKC on Fpn. Our finding that T329E and T361E fail to internalize upon hepcidin-induced ubiquitination implicates that, in certain contexts, ubiquitination alone is not sufficient to drive the endocytosis of Fpn.

### The loss-of-function of PKCα attenuates HH-associated iron overload

HH is a genetic disorder that displays systemic iron overload due to constantly active Fpn (41). Iron overload in multiple organs of HH patients mediates the development of arthritis, diabetes, heart abnormality, liver cirrhosis and cancer (42). Because of the above findings of a stimulatory role of PKC in Fpn expression and iron transport activity, we evaluated whether PKC may be targeted to alleviate body iron overload associated with HH. To address this question, we employed a commonly used HH mouse model Hfe^-/-^ mice, with which Hfe^-/-^;PKCα^-/-^ mice were then generated. Compared to Hfe^-/-^ mice, Hfe^-/-^;PKCα^-/-^ mice of 4 months old started to show a significant decline in serum iron content in both female and male (**Fig. 7A**). Liver iron content was also significant lower in Hfe^-/-^;PKCα^-/-^ mice at 4 (**Fig. 7B**) and 12 (**Fig. 7C**) weeks of age. The decreased liver iron deposition in Hfe^-/-^;PKCα^-/-^ mice was confirmed by *Perl*’s iron staining (**Fig. 7D**). Next, we determined whether the reduced iron loading in Hfe^-/-^;PKCα^-/-^ mice is dependent on the downregulated expression of Fpn. As expected, Fpn expression in the duodenum was markedly decreased in Hfe^-/-^;PKCα^-/-^ mice with a more pronounced decrease in female (**Fig. 7E**), consistent with findings in the WT background (**Fig. 3B**). PKC412 (also called midostaurin) is a pharmacological drug that was initially identified as an inhibitor of PKC, and has been approved for treating acute myeloid leukemia owing to its inhibitory effects on other kinases including FMS-like tyrosine kinase 3 receptor (FLT3) (43, 44). By utilizing PKC412, we determined whether pharmacological inhibition of PKC in *in vivo* could attenuate body iron loading in Hfe^-/-^ mice. To restrict the magnitude of body iron loading prior to PKC412 treatment, newly born Hfe^-/-^ pups along with the dam were put on an iron-deficient diet until the weaning age. We observed that PKC412 treatment for 2 weeks resulted in a significant decrease in duodenal Fpn expression in female and male Hfe^-/-^ mice (**Fig. 7F**). Consistent with the reduction in Fpn expression, serum (**Fig. 7G**) and liver (**Fig. 7H**) iron contents were also significantly decreased in PKC412-treated groups. These results demonstrate that targeting PKC could successfully mitigate Fpn expression and iron overload associated with HH.

**Figure 7.**
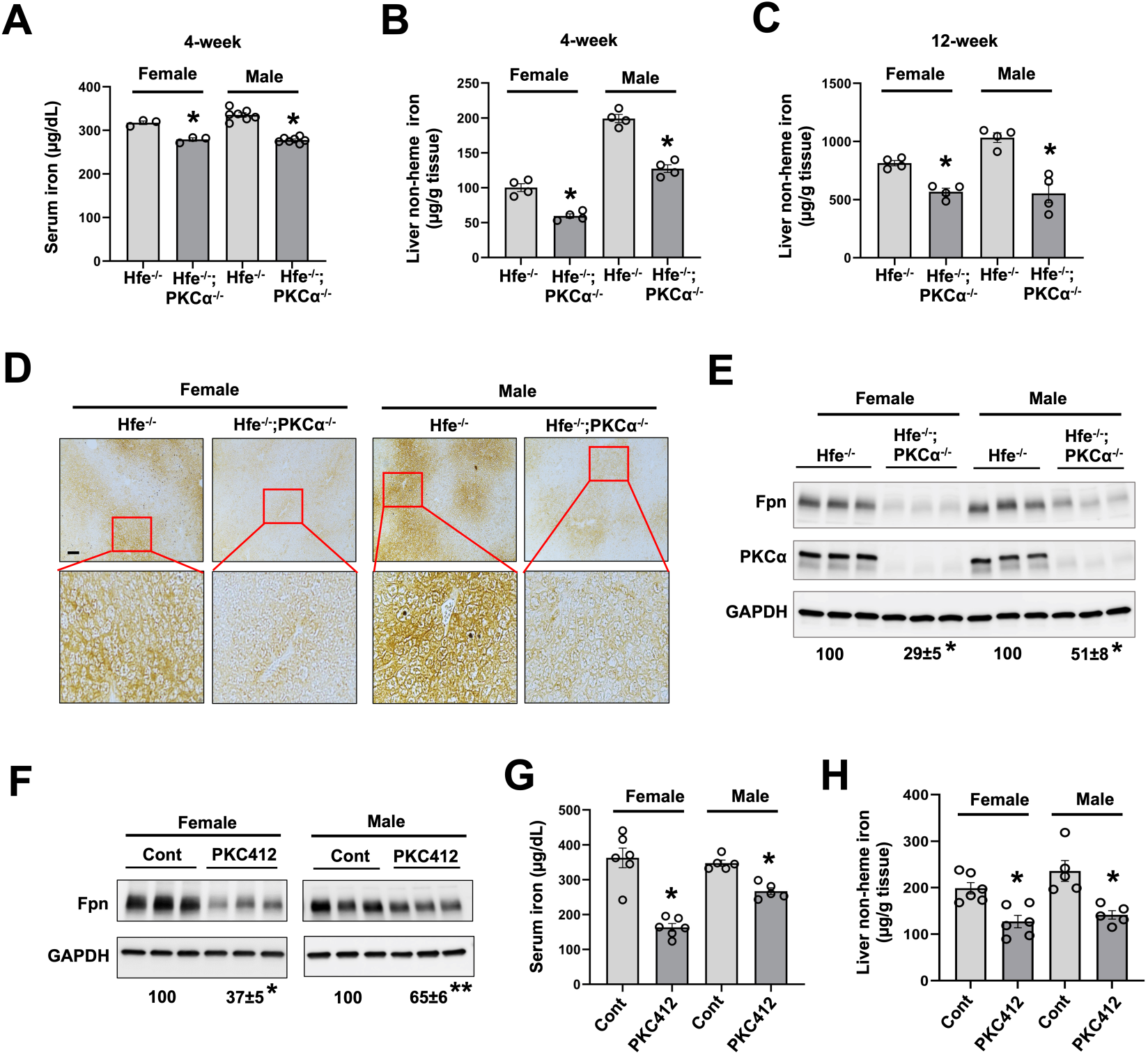
The loss-of-function of PKCα mitigates iron overload in hereditary hemochromatosis. (**A**) Serum iron level was measured in female and male Hfe^-/-^ and Hfe^-/-^; PKCα^-/-^ mice (4 weeks old). Liver non-heme iron content was measured in female and male Hfe^-/-^ and Hfe^-/-^; PKCα^-/-^ mice at 4 (**B**) and 12 (**C**) weeks of age. (**D**) *Perl*’s iron staining shows iron deposition in the liver of female and male Hfe^-/-^ and Hfe^-/-^; PKCα^-/-^ mice. (**E**) Duodenal Fpn expression in female and male Hfe^-/-^ and Hfe^-/-^; PKCα^-/-^ mice by immunoblotting with anti-mouse Fpn antibody. (**F**-**H**) Female and male Hfe^-/-^ mice were fed an iron-deficient diet from when they were born until 3 weeks of age. Mice were then transferred onto chow diet and receive PKC412 (50 mg/kg b.w.) or vehicle for 2 weeks. Duodenal Fpn expression (**F**), serum (**G**) and liver (**H**) iron content were determined. Data are expressed as mean ± SEM (n = 3-7). * *P* < 0.01, and ** *P* < 0.05 compared with Hfe^-/-^ mice or the corresponding controls. *Bar*, 100 μm.

## Discussion

Diabetes is associated with increased body iron stores in humans and experimental models, but the underlying molecular mechanisms remain not well understood. We showed in the current work that Fpn is consistently upregulated in the duodenum accounting for increased iron absorption in multiple mouse models of diabetes and in human patients with type 2 diabetes. For the first time, we have identified that hyperactive PKC signaling associated with hyperglycemia is responsible for the upregulation of Fpn and iron efflux. We showed that PKC upregulates Fpn in both the enterocytes and macrophages, two major cell types that are important for systemic iron homeostasis. In the physiological setting, exocytotic and endocytic trafficking of Fpn are balanced to achieve controlled iron efflux (**Fig. 8, *left***). In the context of hyperglycemia, hyperactive PKC enhances the exocytotic insertion of cytoplasmic Fpn to the plasma membrane (**Fig. 8, *right***). The activation of PKC also attenuates the internalization and degradation of membrane expressed Fpn potentially through the inhibition of hepcidin-mediated ubiquitination of Fpn and increased phosphorylation of Fpn (**Fig. 8, *right***).

**Figure 8.**
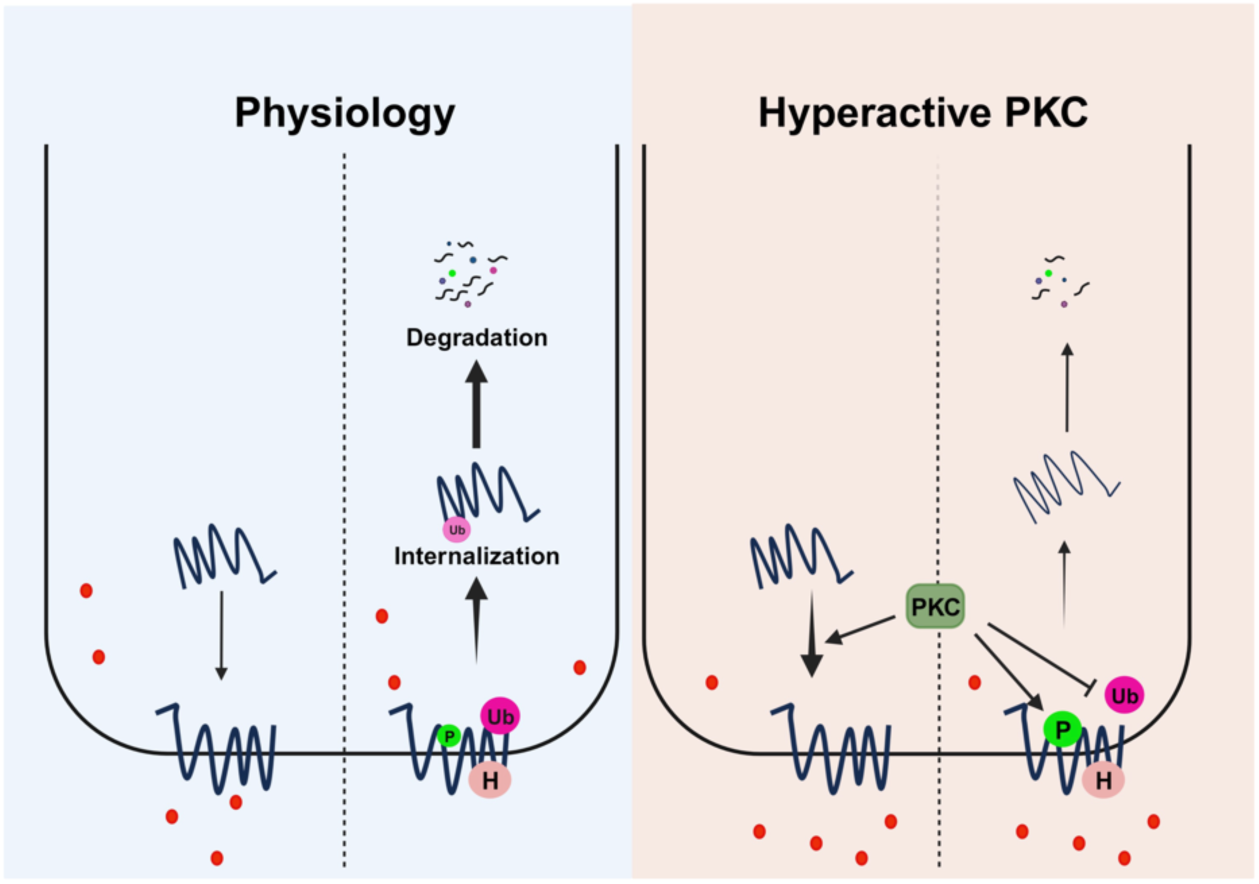
Mechanisms of PKC-promoted membrane expression of Fpn and iron efflux. In the physiological state, the membrane trafficking of Fpn and hepcidin induced ubiquitination-dependent internalization and degradation of Fpn is counterbalanced. In diabetes, wherein the PKC signaling is hyperactivated, the exocytosis of Fpn is enhanced. Hyperactive PKC also increases the phosphorylation and attenuates hepcidin induced ubiquitination of Fpn, which collectively suppress the internalization and degradation of Fpn, leading to increased membrane Fpn expression and iron efflux. *P*, phosphorylation; *H*, hepcidin; *Ub*, ubiquitination; and *red dot* denotes iron.

In this study, to gain a better understanding of the systemic iron status in diabetic context, we employed three mouse models of diabetes including STZ-induced type 1 as well as type 2 diabetic db/db and KKAy mice. Our finding of the increased circulatory and hepatic iron contents in the STZ model is consistent with previous reports (22, 29). The elevation in serum iron in our db/db mice is in line with observations from two other groups (45, 46). We have also observed liver iron loading in the hepatocytes and Kupffer cells in db/db mice. This finding is supported by a previous report of increased liver iron staining in db/db mice (47). However, Altamura *et al.* observed hepatic iron deficiency in db/db mice despite an increase in systemic iron content (48). The discrepancy is unclear, but we noticed a heavy iron loading in hepatic macrophages in our db/db mice. Iron deposition in macrophages is often associated with liver inflammation such as in non-alcoholic steatohepatitis (49, 50). It’s possible that db/db mice with different pathological states display differential patterns of iron deposition. Intriguingly, although KKAy mice manifested a prominent increase in serum iron compared to their controls, like db/db mice in the study by Altamura *et al*. KKAy mice developed a deficiency in liver iron. Our data suggest that decreased expression of hepatic TfR1 might be the underlying mechanism, but it is not understood how TfR1 is downregulated in this genetic background. Regardless, we have shown that Fpn expression in both the duodenum and spleen were consistently upregulated in all three diabetic mouse models. A significant increase in Fpn expression was also seen in the duodenum of type 2 diabetic patients. It is intriguing that multiple anemic diabetic subjects did not show defects in Fpn expression but rather showed a tendency of increased Fpn expression in the duodenum (**Fig. S1** and **Table S4**). Consistent with the upregulation of Fpn, a low intracellular iron level was seen in the enterocytes of diabetic subjects compared to the non-diabetic controls. Previous studies have demonstrated that deficiency in the production of erythropoietin, a kidney-produced hormone that mediates erythropoiesis, is the primary cause of anemia among diabetic patients (51, 52). One would speculate that the increased iron absorption with a decreased iron utilization for red blood cell production could lead to elevated tissue iron overload that aggravates diabetic complications.

Despite the consistent increases in serum iron and Fpn expression, hepcidin expression was divergent in the studied diabetic models. Hepcidin was unaltered in the STZ model, modestly decreased in db/db mice, and mildly increased in KKAy mice, as compared to their corresponding controls. All the three models have a comparable blood glucose level, and therefore it does not seem that the degree of hyperglycemia is a causal factor that influences hepcidin production. Our finding of unchanged hepcidin in STZ model is discrepant from a previous report of decreased hepcidin expression in STZ-induced diabetic rats (22). The study attributed the downregulation of hepcidin to decreased expression of STAT3, a transcription factor that transcribes the expression of hepcidin in inflammation (53, 54). It is unclear whether STAT3 is differentially regulated between diabetic mice and rats. The downregulated expression of hepcidin in db/db mice has previously also been reported (48). However, hepcidin deficiency in db/db mice is likely due to the loss of leptin receptor-mediated signaling. It was shown that leptin induces STAT3-dependent expression of hepcidin in HuH7 human hepatoma cells (36). A later study showed that administration of recombinant leptin to leptin-deficient ob/ob mice led to a significant increase in the mRNA expression of hepcidin (37). In this study, we did not attempt to interrogate how hepcidin is differentially regulated, but the different trends in hepcidin expression in the currently studied models indicates that hepcidin is not the primary factor accounting for Fpn upregulation in the diabetic context.

Many of the PKC family members are activated in hyperglycemia and play important roles in the development of diabetic complications, including diabetic nephropathy, neuropathy, retinopathy, and cardiovascular pathologies (26). Diabetes also provokes changes in the digestive tract to compensate for the systemic deficit in glucose metabolism or utilization. It was reported that intestinal absorption of glucose is increased by multi-folds in STZ-induced diabetic rats, attributable to increased expression of sugar carriers including SGLT1, GLUT2 and GLUT5 (55). Increased GLUT1 expression at the basolateral membrane of the enterocytes was also shown in diabetic rats (56). Studies have shown a potential role of PKC in SGLT1 and GLUT2 regulation by facilitating or stabilizing their expression at the luminal or basolateral membranes in the enterocytes (57). Lee *et al*. reported that PKC phosphorylates GLUT1 at S226 enhancing surface localization of GLUT1 and glucose uptake (58). Iron is a transition metal that plays a crucial role in cellular metabolism dependent on the biosynthesis of iron-sulfur clusters, which are essential cofactors for many enzymes that mediate the citric acid cycle and mitochondrial respiration (59). In the current study, we demonstrated for the first time that the hyperglycemia-PKC axis, *via* the upregulation of Fpn, also promotes the intestinal absorption of dietary iron. We speculate that PKC-dependent activation of iron exporter Fpn, and upregulation of iron importer DMT1 in our previous report (23), may serve as an adaptive mechanism to enhance energy production in diabetes.

Fpn plays a crucial role in iron homeostasis. Much work has focused on its regulation by hepcidin, a negative regulator of Fpn. Herein, we have identified that PKCs, at least the α and 8 isoforms, are novel binding proteins and positive regulators of Fpn. PKCs regulate membrane localization of Fpn by increasing the exocytosis and decreasing the endocytosis of Fpn in the resting and hepcidin treated conditions. However, PKCα did not interfere with the binding of hepcidin to Fpn. Although not determined in our study, it is possible that hyperactive PKC disrupts the ubiquitination process of Fpn. The ubiquitination of Fpn potentially involves E1 enzyme UBA6 (60), and E3 ligase RNF217 (61). It was shown that lysine residues within the region of amino acid 229-269 in the third intracellular loop of the Fpn protein are key sites of ubiquitination (38). One possibility is that PKC binding to Fpn hinders Fpn interaction with the candidate ubiquitination enzymes. It is also possible that PKC signaling interferes with ubiquitination machinery assembly that is required for the ubiquitination of Fpn. These open questions need to be addressed in future studies.

PKCs are serine and threonine protein kinases. As direct interacting proteins of Fpn, PKCs have a high potential of phosphorylating Fpn. *In silico* and mutation analyses led us the finding that PKCs potentially phosphorylate Fpn at T329 and T361, counteracting hepcidin-induced endocytic trafficking of Fpn, despite direct evidence of Fpn phosphorylation by PKCs remains to be confirmed. This phosphorylation dependent stabilization of Fpn does not overlap with PKC-mediated suppression of Fpn ubiquitination, as the phosphomimetic T329E and 361E mutants were equally ubiquitinated like the WT Fpn. It is not understood how the phosphorylation prevents ubiquitinated Fpn from endocytosis. Our recent proteomic analysis identified multiple PDZ domain proteins in the Fpn interactome (unpublished data). PDZ domain proteins are important scaffold proteins that connect many membrane proteins with the cytoskeleton (62). It is tempting to speculate that the phosphorylation of Fpn by PKC strengthens Fpn association with the cytoskeleton preventing the endocytic trafficking of Fpn.

Given the important role of PKCs in Fpn regulation, the loss-of-function of PKCα mitigated diabetes-associated iron overload in diabetic mice. We have further demonstrated that PKC targeting alleviated iron overload in hereditary hemochromatosis by genetic deletion of PKCα and pharmacological inhibition of PKC with PKC412. PKC412 was initially identified as an inhibitor of PKC, and later found that its metabolites can also inhibit multiple other kinases including FLT3, VEGFR-2, JAK3, and RET (63). For its inhibitory effect on FLT3, PKC412 has been used to treat acute myeloid leukemia (AML) (64). Currently, iron overload associated with HH and other disease conditions is managed primarily by phlebotomy or treatment with iron chelators (65). Our study, by demonstrating the efficacy of PKC412 in suppressing Fpn expression and body iron loading, highlights that PKC targeting may be an alternative strategy in the control of iron absorption and recycling to combat iron overload in HH and other pathological conditions.

In conclusion, our study identified for the first time that PKCs are novel interacting proteins and positive regulators of Fpn. Hyperglycemia-mediated activation of PKCs upregulates Fpn-dependent intestinal iron absorption accounts, at least in part, for systemic iron excess in diabetes. PKCs facilitates the membrane expression of Fpn through multiple mechanisms involving the phosphorylation of Fpn, inhibition of hepcidin-dependent and -independent internalization of Fpn, and elevation of exocytotic trafficking of Fpn. A few questions remain to be addressed in future studies including the molecular machinery that mediates PKC-dependent exocytotic and endocytic trafficking of Fpn, the detailed mechanisms by which PKCs suppresses hepcidin-mediated Fpn ubiquitination, and the precise role of PKC-mediated phosphorylation in membrane stabilization of Fpn.

## Methods

### Reagents

The materials and reagents used in the present study are listed in Table S1.

### Animal models

Wild-type (WT) C57BL/6 mice, Hfe^-/-^, db/+, db/db, KKAy, and the a/a control mice were purchased from the Jackson Laboratory. PKCα^-/-^ mice in C57BL/6 background were as previously described (23), and were used for crossbreeding with db/+ and Hfe^-/-^ mice to obtain db/db;PKCα^-/-^ and Hfe^-/-^;PKCα^-/-^ mice, respectively. To induce type 1 diabetes, mice at the age of 6-8 weeks were given streptozotocin (STZ, 40 mg/kg body weight) for five consecutive days. Mice with fasting blood glucose levels above 300 mg/dl were selected for the study. STZ-treated, db/db and KKAy mice were sacrificed approximately two months after they were on hyperglycemia, unless otherwise noted. For the study involving PKC412, the newly born Hfe^-/-^ pups with their dams were fed an iron-deficient diet (TD.80396) till weaning age. After weaning, mice were transferred onto a regular diet and received PKC412 (50 mg/kg/day, 200 μl DMSO:PEG300:H_2_O in 5:45:50 ratio) *via* oral gavage for two weeks. At the end of PKC412 treatment mice were sacrificed, and samples were collected for the subsequent analyses.

### Site-directed mutagenesis

The phosphorylation sites in mouse and human Fpn proteins were predicted by employing multiple phosphorylation site prediction online tools including NetPhos 3.1, NetPhorest, and PhosphoMotif Finder. The phospho-dead and phosphomimetic mutants of pEGFP/hFpn were generated by site-directed mutagenesis using QuikChange II XL Site-Directed Mutagenesis kit following the manufacturer’s instructions. Mutagenesis primers were synthesized by the Integrated DNA Technologies, and the primer sequences are listed in Table S2. All the mutant constructs were verified through sequencing by Eurofins Genomics.

### Cell Culture and Transfection

Human renal proximal tubular epithelial HEK293T from ATCC were cultured in DMEM (high glucose) supplemented with 10% fetal bovine serum, 50 U/ml penicillin, and 50 μg/ml streptomycin. HEK293T cells were transfected with pEGFP/hFpn, phospho-dead or phosphomimetic Fpn mutants, by using Lipofectamine 2000. Dox inducible TRex cell line was as previously described (38). The expression of Fpn-GFP was induced either with 1 µg/ml Dox for up to 4-h incubation or with 100 ng/ml Dox for overnight induction. BMDMs were isolated from the femur and tibia bones of 6-8 weeks old mice and cultured in DMEM supplemented with 10% heat-inactivated FCS, 2% HEPES, 1% non-essential amino acid, 1% sodium pyruvate, 1x antibiotic/antimycotic, and macrophage colony-stimulating factor (M-CSF, 10 ng/ml, added every other day) in 6-well plates. After 6 days of culture, the cells were washed twice with DMEM to remove floating cells, and then cultured in fresh medium excluding M-CSF for 24 h prior to any treatments.

### Quantitative reverse transcription PCR (qRT-PCR) analysis

Mouse livers were stored in RNAlater and homogenized for total RNA extraction using RNeasy® Plus Mini Kit. Two µg of total RNA were used for cDNA synthesis by using the High-capacity cDNA Reverse Transcription Kit. Quantitative PCR was performed using Applied Biosystems SYBR PCR Master Mix on QuantStudio^TM^ 3 Real-Time PCR System. The primers used for qPCR are included in Table S3. The relative expression of the target gene was calculated using the comparative 2^-ΔΔCt^ method (66).

### Western blotting

The duodenum (the first 3 cm of the intestine) was dissected, longitudinally cut, and washed for 5 min in cold phosphate-buffered saline (PBS). Crude villus debris was obtained through a 30-min incubation in cold PBS containing 10 mM EDTA, followed by a gentle vortex for 15 sec. The isolated villi and spleen were homogenized and sonicated in RIPA lysis buffer supplemented with protease and phosphatase inhibitor cocktail. Protein supernatants were obtained following a 10-min centrifugation at 14, 000x g, and the concentration was determined by the BCA method. Protein lysates were incubated in 1x Laemmli buffer for 30 min at room temperature (RT). Proteins were then separated on SDS-PAGE gel and transferred onto nitrocellulose membrane (0.45 µm) for immunoblotting with the corresponding antibodies. Densitometric analysis was performed with the ImageJ software (NIH, Bethesda, MD).

### Immunoprecipitation

The cells were washed twice in cold PBS and lysed in the Cell Lysis Buffer. The collected crude lysate was sonicated twice for 10 sec and centrifuged at 14 000x g for 10 min. Lysates were incubated with the anti-rabbit GFP antibody for 4 h, and then incubated for 2 h with protein A/G magnetic beads, followed by three washes in cold PBS containing 1% Triton X-100 and a final wash with PBS only. All procedures were performed at 4 °C or on ice. The bound immunocomplex was eluted by incubating the beads for 7 min at 95 °C in 2X Laemmli buffer supplemented with 2.5% β-Mercaptoethanol, followed by SDS-PAGE separation and Western blot analysis.

### Surface biotinylation

Cell surface biotinylation was performed as in our previous study (67). In brief, BMDMs and 293T cells were washed once in cold PBS and incubated for 10 min in borate buffer (154 mM NaCl, 7.2 mM KCl, 1.8 mM CaCl_2_, and 10 mM H_3_BO_3_, pH 9.0). Then the cells were incubated for 40 min with 0.5 mg/ml NHS-SS-biotinin in borate buffer. Unbound biotin NHS-SS-biotin was quenched by incubating the cells with Tris buffer (20 mM Tris, 120 mM NaCl, pH 7.4) for 5 min. Cells were rinsed with PBS and lysed in the Cell Lysis Buffer. An aliquot of the lysate was retained as the total fraction. Approximately 200-400 µg of lysate was incubated with streptavidin-agarose beads for 2 h. Agarose beads were then washed, and bound immunocomplex was eluted as in the above described.

### Hepcidin binding assay

Hepcidin and Fpn binding assay was performed as previously described (68). Briefly, cells were treated with 3 µg/ml Biotinyl-Hepcidin-25 for 30 min at 37 °C, followed by lysis in the Cell Lysis Buffer. The protein lysates were subjected to immunoprecipitation with anti-rabbit GFP antibody. The immunocomplex was boiled for 7 min in 2x Laemmli buffer in the absence of reducing reagent. The eluted proteins were analyzed by immunoblotting with HRP-conjugated streptavidin.

### Immunofluorescence staining

Freshly isolated mouse duodenum was fixed in 4% paraformaldehyde for 60 min, and then incubated in 30% sucrose overnight at 4 °C. Duodenum was embedded in OCT and cryosections were prepared. Human duodenal biopsy was fixed in 10% formalin and paraffin sections were made. Formalin-fixed paraffin-embedded (FFPE) duodenal sections from type 2 diabetic (n = 14) and non-diabetic control (n = 14) human subjects were obtained from the pathology archive at Brown University. The detailed demographic and clinical information of the human subjects is summarized in Table S4. FFPE sections were rehydrated and subjected to antigen retrieval in citrate buffer (10 mM, pH 6.0). The above processed tissue sections were permeabilized in PBS containing 0.2% Triton-X100, followed by blocking with 5% normal goat serum. The specimen was then incubated for 1 h with the corresponding primary antibodies at RT. After washing, the sections were incubated for 30 min with Alexa Fluor-conjugated secondary antibodies and DAPI. The sections were mounted with ProLong Gold Antifade mounting medium and visualized under an Olympus 3x FV1000 confocal or Leica DM6 B widefield fluorescence microscope.

### Liver and serum iron measurement

To measure liver iron content, the same weight of wet liver tissue was homogenized in a protein precipitation solution (0.53 g Trichloroacetic acid and 440 µl of HCl in 10 ml H_2_O) (69). The suspension was then incubated for 10 min at RT and boiled for 1 h. The suspension was cooled to RT and centrifuged at 14,000x g for 10 min to collect the supernatant. To prepare serum, blood collected from mice was seated at RT for 30 min without disturbance, followed by a centrifugation at 2,500x g for 20 min. The obtained liver lysate and serum were used for the measurement of non-heme iron content using the Iron Assay kit.

### Iron staining

Iron content in the paraffin-embedded liver sections of mice was determined by *Prussian* blue iron staining (31). In brief, the tissue sections were washed in distilled water after being deparaffinized and rehydrated in graded series of xylene and ethanol. Then, the sections were incubated for 30 min in 5% potassium ferrocyanide and 10% hydrochloric acid solution. For enhanced *Perl*’s iron staining, tissue sections were further incubated for 20 min in 0.3% (v/v) hydrogen peroxide diluted in methanol, followed by incubation in 0.05% diamine benzidine tetrahydrochloride (DAB) and 0.01% hydrogen peroxide in distilled water. The sections were rapidly dehydrated in ethanol and xylene, mounted, and imaged using Leica DMC4500 microscope.

### Serum hepcidin ELISA

Mouse serum hepcidin level was measured using a hepcidin (mouse) ELISA kit following the manufacturer’s instruction.

### Data availability

All data obtained throughout the study are included within this article and supplementary files.

### Statistical analysis

Unpaired *t*-test and one-way analysis of variance (ANOVA) test were employed to determine the statistical significance using GraphPad Prism 9.0 software. Data are presented as Mean ± SEM, and *p* < 0.05 was considered statistically significant.

### Study approval

All the animal experiments were performed under approval by the Institutional Animal Care and Use Committee of Emory University and in accordance with the US National Institutes of Health Guide for the Care and Use of Laboratory Animals. Research with the de-identified human samples was approved by the Human Ethics Committee of Brown University and performed in accordance with the NIH and institutional guidance for human subject research.

## Author contributions

SB, HT, and PH conducted the experiments; SL screened and provided human duodenal samples; SB and IW performed data analysis; SB, EN, and PH designed the experiments and wrote the manuscript; and SL, AJ, SS, EN, and PH edited the manuscript.

## Acknowledgements

This study was supported by the National Institute of Health grant R01DK125647 (to PH). This work utilized the Emory University Integrated Cellular Imaging Core Facility that is supported by RRID: SC_023534.

