## Supplemental figures and tables for "Targeting PKC alleviates iron overload in diabetes and hemochromatosis"

### Supplementary materials

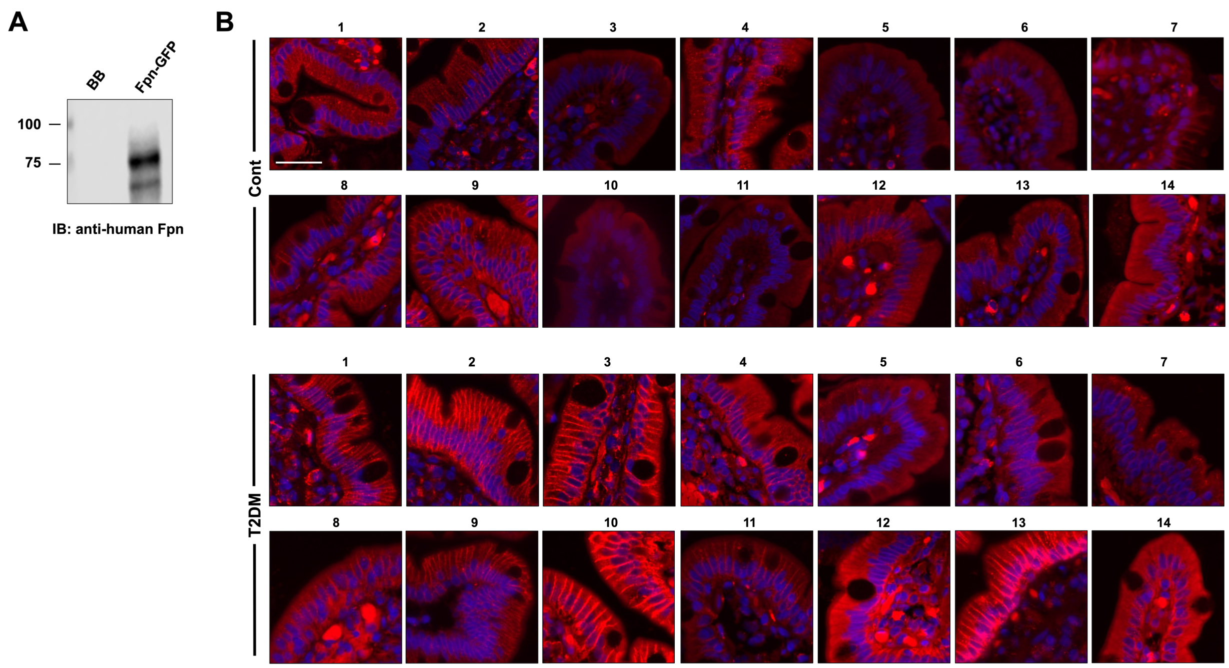


**Fig. S1. Increased Fpn expression in the duodenum of human diabetic patients**. (**A**) Validation of the rabbit anti-human Fpn antibody (R&D) by using 293T lysates with and without the expression of Fpn-GFP. (**B**) Fpn expression in the duodenal samples of type 2 diabetic (T2DM, n = 14) and non-diabetic control (n = 14) subjects with anti-human Fpn antibody, followed by fluorescent imaging. DAPI was used to counterstain the nuclei. *Bar*, 50 μm.

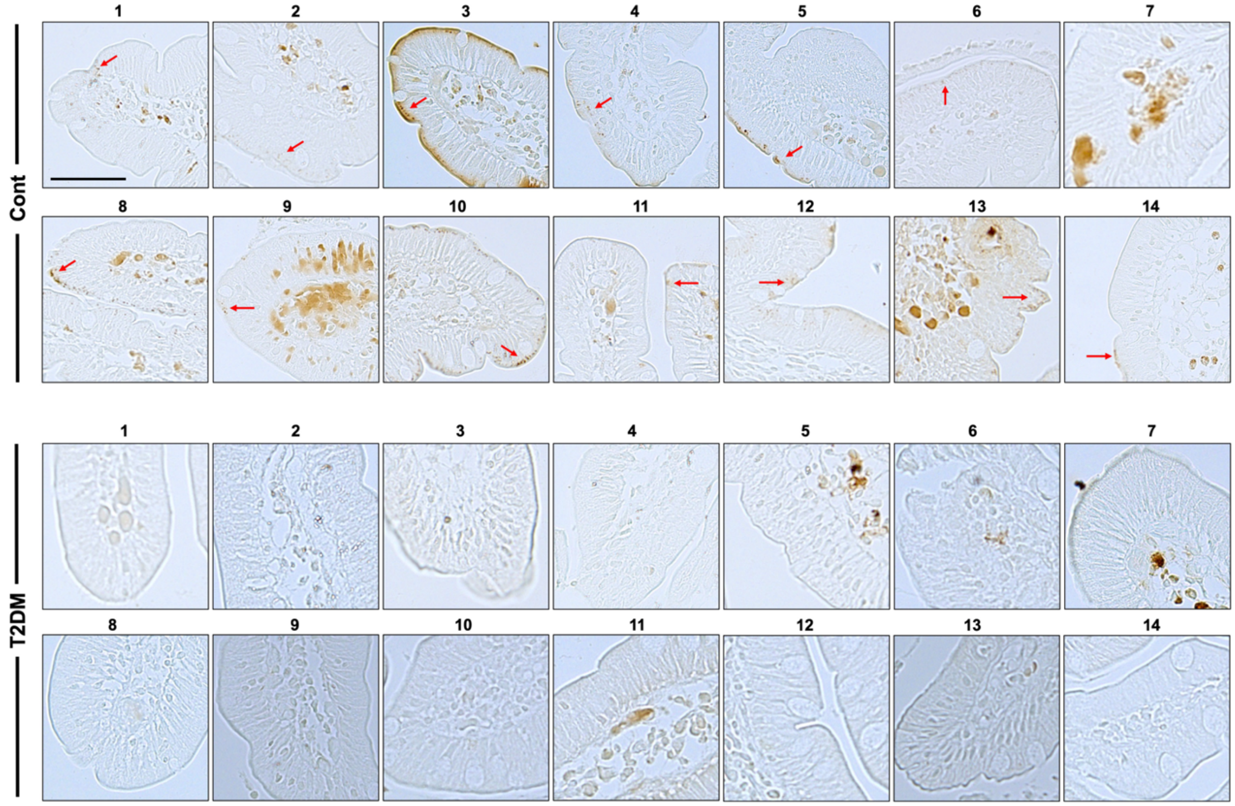


**Fig. S2. Iron content is decreased in the enterocytes of diabetic patients**. *Perl*'s iron staining was performed to detect iron content in duodenal enterocytes of type 2 diabetic (T2DM, n = 14) and non-diabetic control (n = 14) subjects. *Red arrow* denotes epithelial iron deposition. *Bar*, 50 μm.

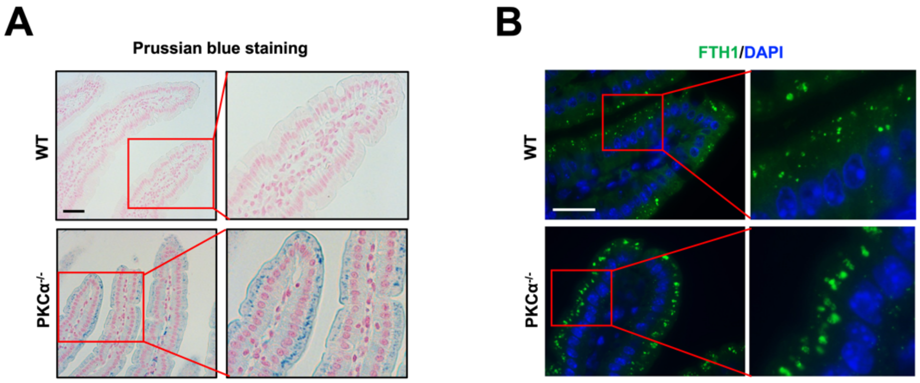


**Fig. S3. The loss-of-function of PKCα induces intestinal epithelial iron accumulation**. Iron sequestration in duodenal epithelium was determined by *Prussian* blue staining (**A**) and staining for ferritin heavy chain 1 (FTH1) (**B**) in 16-week-old WT and PKCα$^{-/-}$ mice. Nuclei was stained with DAPI. *Bar*, 50 μm.

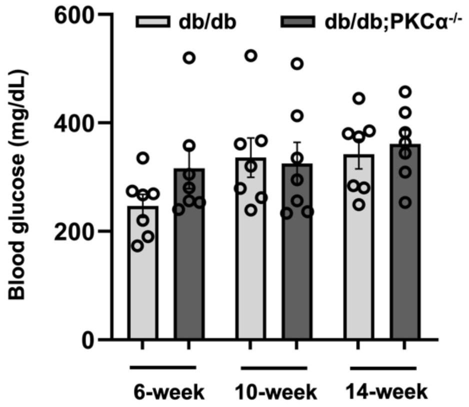


**Fig. S4. Knockout of PKCα does not alter blood glucose level in db/db mice**. Blood glucose was monitored in db/db and db/db;PKCα$^{-/-}$ mice (female and male) starting from 6 weeks to 14 weeks of age. Data shown are mean ± SEM (n = 7).

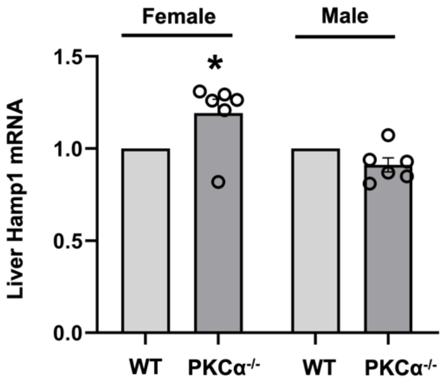


**Fig. S5. The effect of the knockout of PKCα on liver hepcidin expression**. Liver mRNA expression was determined by qRT-PCR in 16 weeks old female and male WT and PKCα$^{-/-}$ mice. Data shown are mean ± SEM (n = 6). * *P* < 0.01 compared to the WT mice.

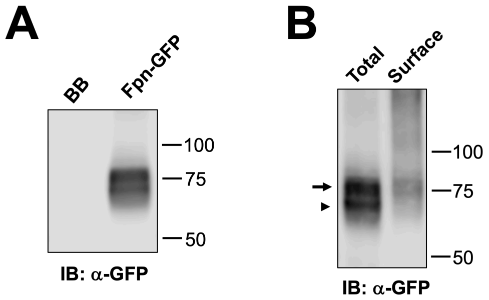


**Fig. S6. Size difference in cytoplasmic and membranous Fpn.** (**A**) Fpn-GFP protein presented in two bands in 293T cells by Western blotting with anti-GFP antibody. (**B**) Fpn-GFP protein in the total homogenates and membrane pool differed in band pattern. *Arrow*, membrane form; *arrowhead*, cytoplasmic form.

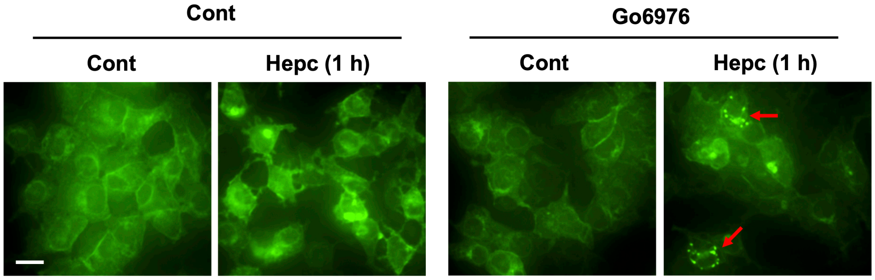


**Fig. S7. Inhibition of PKCα sensitizes Fpn to hepcidin-induced internalization**. 293T cells transfected with Fpn-GFP were pretreated for 1 h with 2 μM Go6976. Cells were then incubated for 1 h with 0.5 μg/ml hepcidin (Hepc) prior to florescent imaging. *Red arrow* denotes cytoplasmic Fpn-GFP. *Bar*, 10 μm.

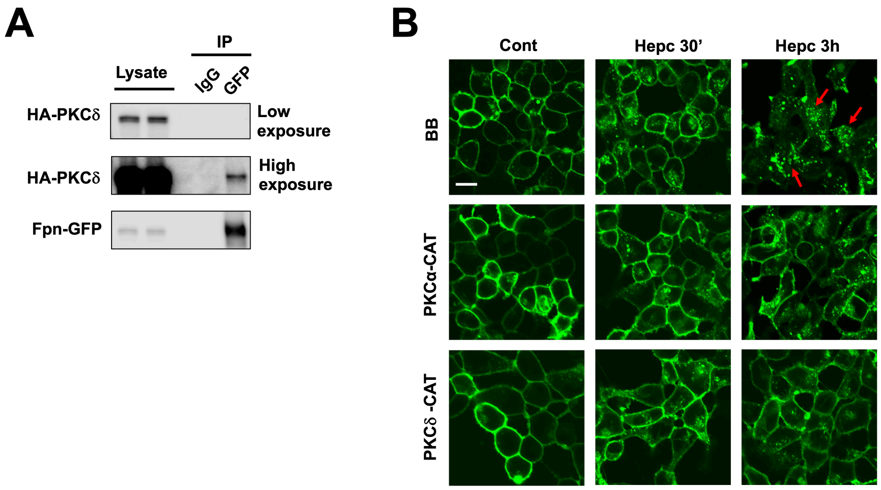


**Fig. S8. PKCδ binds and suppresses hepcidin-induced internalization of Fpn**. (**A**) Fpn-GFP was immunoprecipitated from Dox-treated TRex cells that were transfected with full-length HA-PKCδ. The presence of HA-PKCδ in the immunocomplex was determined by blotting with anti-HA antibody. (**B**) TRex cells transfected with vector backbone (BB), PKCα-CAT, or PKCδ-CAT were induced with Dox prior to treatment with or not 0.5 μg/ml hepcidin for 30 min or 3 h. The subcellular localization of Fpn-GFP was then determined by confocal microscopy. *Red arrow* denotes cytoplasmic Fpn-GFP. *Bar*, 10 μm.

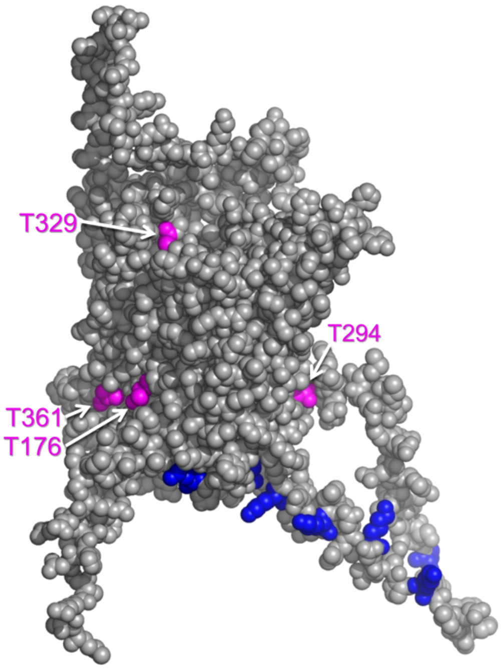


**Fig. S9. The location of the predicated PKC phosphorylation sites on the 3D structure of human Fpn**. PDB file for human Fpn was obtained from AlphaFold protein structure database. T176, T294, T329 and T361 are colored in magenta. Lysines in the large intracellular loop that are implicated as targets of ubiquitination after hepcidin binding are colored in blue (K229, 236, 240, 247, 253, 258, 269).

**Table S1**. **List of reagents, antibodies, and plasmid constructs.**

| | Source | Catalog | Species | Description |
|---|---|---|---|---|
| **Reagents** | | | | |
| 4X Laemmli Sample Buffer | Bio-Rad | 1610747 | | |
| Antibiotic:Antimycotic Solution | Gemini Bio Products | 400101 | | |
| Bicinchoninic Acid Assay kit | Thermo Fisher Scientific | 23225 | | |
| Biotinyl Hepcidin | Bachem | 4056950 | | |
| Boric acid | Sigma-Aldrich | B6768 | | |
| Cell Lysis Buffer (10x) | Cell Signaling Technology | 9803 | | |
| Citrate buffer solution 0.09 M | Sigma-Aldrich | C2488 | | |
| Cycloheximide | Sigma-Aldrich | C7698 | | |
| DAPI | Cell Signaling Technology | 4083 | | |
| DMEM, High Glucose | Gibco | 11960-044 | | |
| Doxycycline hydrochloride | Sigma-Aldrich | D3447 | | |
| EZ-LinkTM Sulfo-NHS-SS-Biotin | Thermo Fisher Scientific | 21331 | | |
| Fetal Bovine Serum | R&D Systems | S11150 | | |
| GlutaMAX$^{TM}$ Supplement | Gibco | 35050061 | | |
| Go6976 | Cell Signaling Technology | 12060 | | |
| Go6983 | Tocris | 2285 | | |
| HaltTM Protease and Phosphatase Inhibitor Cocktail (100X) | Thermo Fisher Scientific | 1861282 | | |
| Hepcidin (Human) | Peptide Institute, Inc. | 4392-s | | |
| Hepcidin (Mouse) | Peptide Institute, Inc. | 4434-s | | |
| Hepcidin (mouse) ELISA kit | BioVision | E4693 | | |
| High Capacity Streptavidin Agarose | Thermo Fisher Scientific | 20357 | | |
| High-Capacity cDNA Reverse Transcription Kit | Applied Biosystems | 4368814 | | |
| Iron Assay Kit | Sigma-Aldrich | MAK025 | | |
| Iron Deficient Diet | Envigo | TD.80396 | | |
| Lipofectamine 2000 | Thermo Fisher Scientific | 11668-019 | | |
| PEG300 | Spectrum Chemical | PO108 | | |
| PKC412 | Combi-Blocks | HE-3861 | | |
| Potassium chloride | Sigma-Aldrich | P4504 | | |
| Potassium hexacyanoferrate (II) trihydrate | Sigma-Aldrich | P3289 | | |
| Prolong Diamond Antifade Mounting Medium | Thermo Fisher Scientific | P36961 | | |
| Protein A/G Magnetic Beads | Thermo Fischer Scientific | 88802 | | |
| QuikChange II XL Site-Directed Mutagenesis Kit | Agilent Technologies | 200521 | | |
| Recombinant Mouse M-CSF | R&D Systems | 416-ML | | |
| RIPA Buffer (10X) | Cell Signaling Technology | 9806 | | |
| RNAlater$^{TM}$ Stabilization Solution | Invitrogen | AM7020 | | |

| | | | | |
|---|---|---|---|---|
| RNeasy® Plus Mini Kit | Qiagen | 74136 | | |
| SignalStain® DAB Substrate Kit | Cell Signaling Technology | 8059 | | |
| SsoAdvanced Universal SYBR Green Mastermix | Bio-Rad | 1725271 | | |
| Streptozotocin | Cayman Chemicals | 13104 | | |
| Trichloroacetic acid | Sigma-Aldrich | T6399 | | |
| **Antibodies** | | | | |
| anti-mouse Alexa Fluor 568 | Thermo Fisher Scientific | A11031 | Goat | |
| anti-mouse HRP | Cell Signaling Technology | 7076 | Goat | |
| anti-rabbit Alexa Fluor 488 | Thermo Fisher Scientific | A11034 | Goat | |
| anti-rabbit HRP | Cell Signaling Technology | 7074 | Goat | |
| anti-rat Alexa Fluor 568 | Thermo Fisher Scientific | A11077 | Goat | |
| c-Myc | Sigma-Aldrich | M4439 | Mouse | 1:1000 |
| F4/80 | Cell Signaling Technology | 70076 | Rabbit | 1:1000 |
| Ferroportin | Alpha Diagnostics | MTP11-A | Rabbit | 1:1000 (mouse, WB) |
| Ferroportin | R&D Systems | MAB9924 | Rabbit | 1:1000 (human, IF) |
| FK2 | Cayman Chemical Company | 14220 | Mouse | 1:1000 |
| FTH1 | Cell Signaling Technology | 3998 | Rabbit | 1:1000 |
| GAPDH | Cell Signaling Technology | 5174 | Rabbit | 1:10000 |
| GFP | abcam | ab290 | Rabbit | 1:5000 |
| GFP | Roche | 11814460001 | Mouse | 1:1000 |
| HA-Tag (C29F4) | Cell Signaling Technology | 3724 | Mouse | 1:1000 |
| HRP-Conjugated Streptavidin | Thermo Fisher Scientific | N100 | | 1:1000 |
| PKCα | BD Biosciences | 610108 | Mouse | 1:1000 (IF) |
| PKCα | Cell Signaling Technology | 2056 | Rabbit | 1:1000 (WB) |
| **Plasmids** | | | | |
| N3-hFpn-GFP | Dr. Elizabeta Nemeth lab | | Human | |
| PKC alpha WT | Addgene | 21232 | Human | |
| PKC alpha CAT | Addgene | 21234 | Human | |
| PKC delta WT | Addgene | 16386 | Human | |
| PKC delta CAT | Addgene | 16388 | Human | |

**Table S2. Primers for site-directed mutagenesis of human Fpn-GFP.**

| Mutant | | Primer sequence, 5'→3' |
|---|---|---|
| T176A | For | AACTAGCAAATATGAATGCCGCAATACGAAGGATTGACCAG |
| | Rev | CTGGTCAATCCTTCGTATTGCGGCATTCATATTTGCTAGTT |
| T294A | For | GCTGAGCCCTTCCGTGCCTTCCGAGATG |
| | Rev | CATCTCGGAAGGCACGGAAGGGCTCAGC |
| T329A | For | GTGTAGGCGTACCCTGCGGTGATGCAGTCAAAG |
| | Rev | CTTTGACTGCATCACCGCAGGGTACGCCTACAC |
| T361A | For | GAATAATGGGAACTGTAGCTTTTGCTTGGCTACGTCGAAAAT |
| | Re | ATTTTCGACGTAGCCAAGCAAAAGCTACAGTTCCCATTATTC |
| T329E | For | GAGTGTAGGCGTACCCTTCGGTGATGCAGTCAAAGC |
| | Rev | GCTTTGACTGCATCACCGAAGGGTACGCCTACACTC |
| T361E | For | CATTTTCGACGTAGCCATTCAAAAGCTACAGTTCCCATTATTCCAGTTATAGC |
| | Rev | GCTATAACTGGAATAATGGGAACTGTAGCTTTTGAATGGCTACGTCGAAAATG |

A, Alanine; E, Glutamic acid; and T, Threonine.

**Table S3. Primers for qRT-PCR.**

| **Gene** | **Primer sequence, 5'→3'** | |
|---|---|---|
| | For | Rev |
| **Hamp1** | TTGCGATACCAATGCAGAAGA | GATGTGGCTCTAGGCTATGTT |
| **TfR1** | AATTGGGTGTTGGGAAGACAA | ACATTCTCAGGTGGCAGCTT |
| **ZIP14** | CTGGCTATTGGTGCCTCCTTCA | TGCCAGCATTGAGCAGGATGAC |
| **Rpl13a** | ATGACAAGAAAAAGCGGATG | CTTTTCTGCCTGTTTCCGTA |

**Table S4. Demographic and clinical information of the studied human subjects.**

| ID | | Sex | Age (years) | HbA1C (%)/ Glucose (mg/dL) | Hb (g/dL) |
|---|---|---|---|---|---|
| Control | 1 | F | 22 | Glucose 90 | 13.9 |
| | 2 | F | 42 | 5.2 | 13.4 |
| | 3 | F | 30 | No reading | 13.5 |
| | 4 | F | 57 | 5.2 | 14.2 |
| | 5 | M | 36 | Glucose 91 | 14.9 |
| | 6 | F | 34 | Glucose 86 | 13.6 |
| | 7 | M | 47 | 5.6 | 15 |
| | 8 | F | 33 | Glucose 90 | 14.3 |
| | 9 | F | 25 | No reading | 13.6 |
| | 10 | M | 62 | Glucose 99 | 12.7 |
| | 11 | F | 73 | Glucose 84 | 13.6 |
| | 12 | F | 37 | 5.3 | 12.8 |
| | 13 | F | 18 | No reading | 13 |
| | 14 | F | 56 | 5.6 | 12.7 |
| T2DM | 1 | M | 59 | 7.1 | 13 |
| | 2 | F | 60 | 7.1 | 10.5 |
| | 3 | M | 67 | 7.3 | 10.2 |
| | 4 | F | 58 | 7.1 | 10.4 |
| | 5 | F | 58 | 7.8 | 13 |
| | 6 | F | 55 | 8.5 | 10.9 |
| | 7 | F | 58 | 8.6 | 13.7 |
| | 8 | F | 57 | 7.4 | 12.8 |
| | 9 | F | 70 | 7.8 | 10.1 |
| | 10 | F | 39 | 7.6 | 12.4 |
| | 11 | F | 36 | 7.2 | 13.2 |
| | 12 | M | 69 | 9.4 | 14.1 |
| | 13 | F | 71 | 12.6 | 11.3 |
| | 14 | F | 66 | 7.6 | 12.2 |

F, Female; M, Male; Hb, Hemoglobin; and T2DM, type 2 diabetes mellitus.
